# Building an RNA switch-based selection system for enzyme evolution in yeast

**DOI:** 10.1101/2022.06.14.496130

**Authors:** Deze Kong, Christina Smolke

**Affiliations:** Department of Bioengineering, Stanford University, Stanford, CA, USA; Chan Zuckerberg Biohub, San Francisco, CA, USA

## Abstract

Recent advances in synthetic biology and metabolic engineering have enabled yeast as a favorable platform for synthesis of valuable natural and semi-synthetic compounds through expression of multiple heterologous enzymes sourced from plants, fungi and bacteria. However, these heterologous enzymes can suffer from low activity, specificity, stability and solubility in yeast, resulting in arduous iterations of design-build-test-learn cycles to optimize their production often performed on a single enzyme basis. Laboratory directed evolution has proven to be a powerful and high-throughput method for protein engineering, albeit its limited application for biosynthetic enzymes. Here, we harness small molecule-sensing, RNA-based switches to develop a generalizable selection system facilitating enzyme evolution. Our design utilizes an RNA-based switch for detection of intracellular compound production, which then regulates the expression of a selection gene. Our initial data shows that the auxotrophy selection gene *SpHIS5* exhibits the highest selective capability in combination with a theophylline-responsive RNA-based switch. Using the theophylline-responsive RNA-based switch, we demonstrated the enrichment of a high-producing variant of caffeine demethylase, in a population size of 10^3^. We target to demonstrate the use of this RNA-based selection system as a general approach for enzyme evolution.

## INTRODUCTION

Yeasts provide a powerful platform for design of microbial factories. Previous research demonstrated the incorporation of complex biosynthetic pathways from medicinal plant sources into yeast for production of a variety of valuable pharmaceutical compounds, including the antimalarial drug precursor artemisinic acid, the painkiller opiates, the potential anti-cancer drug noscapine, and the neurological disorder-treating drug tropane alkaloids^1–4^. To achieve enhanced production of complex drug molecules in yeast, the reconstructed biosynthetic pathways for producing the target compounds often undergo various strategies of optimization^5^. Laboratory directed evolution has proven to be an efficient method for enzyme engineering, improving its activity, specificity, or expression in microbial hosts by identifying desired variants from large randomized or designed libraries^6^.

Recent research has explored the application of biosensors for engineering biosynthetic pathways in microbial systems^7^. Specifically, biosensors have been used to detect the intracellular presence of a target compound or metabolite, and thereby regulate the expression of a reporter gene for screen or selection of high-producing strains. RNA-based switches are one category of biosensors which enables gene regulation either through monitoring translation or stabilizing mRNA transcript of a target gene upon ligand binding. Artificial RNA-based switches are designed in a modular manner and consist of a sensor domain, i.e., an RNA aptamer for ligand binding, an actuator domain, i.e., a self-cleaving hammerhead ribozyme for post-transcriptional regulation, and a transmitter domain, i.e., an RNA sequence joining the sensor and the actuator domains and mediating the switch function, designed either rationally or screened *in vitro*^8^.

Here, we target to demonstrate the usage of RNA-based switches as biosensors to correlate yeast intracellular metabolite levels to cellular growth by using an RNA-based switch to regulate the expression of a survival gene, therefore developing a selection system for evolving high-producing enzyme variants. A previous work demonstrated using an RNA-based switch for detecting intracellular production of metabolites in yeast, thus generating a signal for enzymatic activities through regulation of the expression of a reporter gene, i.e., green fluorescent protein gene, for establishing a FACS-based screen^9^. In this work, an enzyme library size of 10^3^ was screened using a theophylline-responsive RNA switch and the enzyme activity was improved by 33-fold, and selectivity by 22-fold *in vivo*.

Other examples demonstrated using RNA-based switches to regulate the expression of a survival or lethal gene, thus establishing a selection system. For example, a previous work used a self-cleaving ribozyme glmS, stabilized by intracellular gluocosamine 6-phospate (GlcN6P), for evolution of a yeast strain producing *N*-acetyl glucosamine (GlcNAc)^10^. The glmS ribozyme was used to regulate the expression of a lethal gene, i.e., *FCY1*, which encodes cytosine deaminase, an enzyme converting fluorocytosine to fluorouracil, a toxin to eukaryotic system. The study demonstrated evolution of two enzymes, a glutamine-fructose-6-phosphate transaminase and a heterologous haloacid dehalogenase-like phosphate using the self-cleaving ribozyme. In another study, a lysine-responsive RNA switch was used to design a dual selection system in *E. coli*, which regulates the expression of a *TetA* selection gene encoding a tetracycline / H^+^ antiporter^11^. In the positive selection mode, activated lysine-producing variants are selected in the presence of the ligand and the selection pressure, tetracycline. In the negative selection mode, inactivated variants are selected in the absence of the ligand and presence of the negative selection pressure, nickel chloride, as expression of TetA leads to increased cellular permeability to toxic nickel salts. In this study, the pathway-optimized lysine-producing strains were enriched up to 75% of the total population after four rounds of enrichment cycles.

Here, we present a pipeline for developing an RNA switch-based selection system in *S. cerevisiae*. Specifically, we use an RNA switch to regulate the expression of an auxotrophic gene or an antibiotic resistance gene to monitor yeast growth (Figure 1). A genetically diversified enzyme library is first transformed into the yeast cell on a low-copy plasmid vector. Each variant from the library then encodes an enzyme, either several steps away, or immediately upstream of the metabolite to be detected by the RNA-based sensor. High activity of the enzyme variant will result in an accumulation of the target metabolite, which will bind to the RNA aptamer, thereby stabilizing its structure and leading to translation of a selection gene. To maximize the dynamic range in growth rates titrated by an RNA switch, we first use the RNA switch to regulate the expression of a transactivator gene, which will then be translated into a transactivator protein. The transactivator protein binds to a transactivator-induced promoter, which activates the expression of the selection gene, leading to yeast growth regulation.

**Figure 1:**
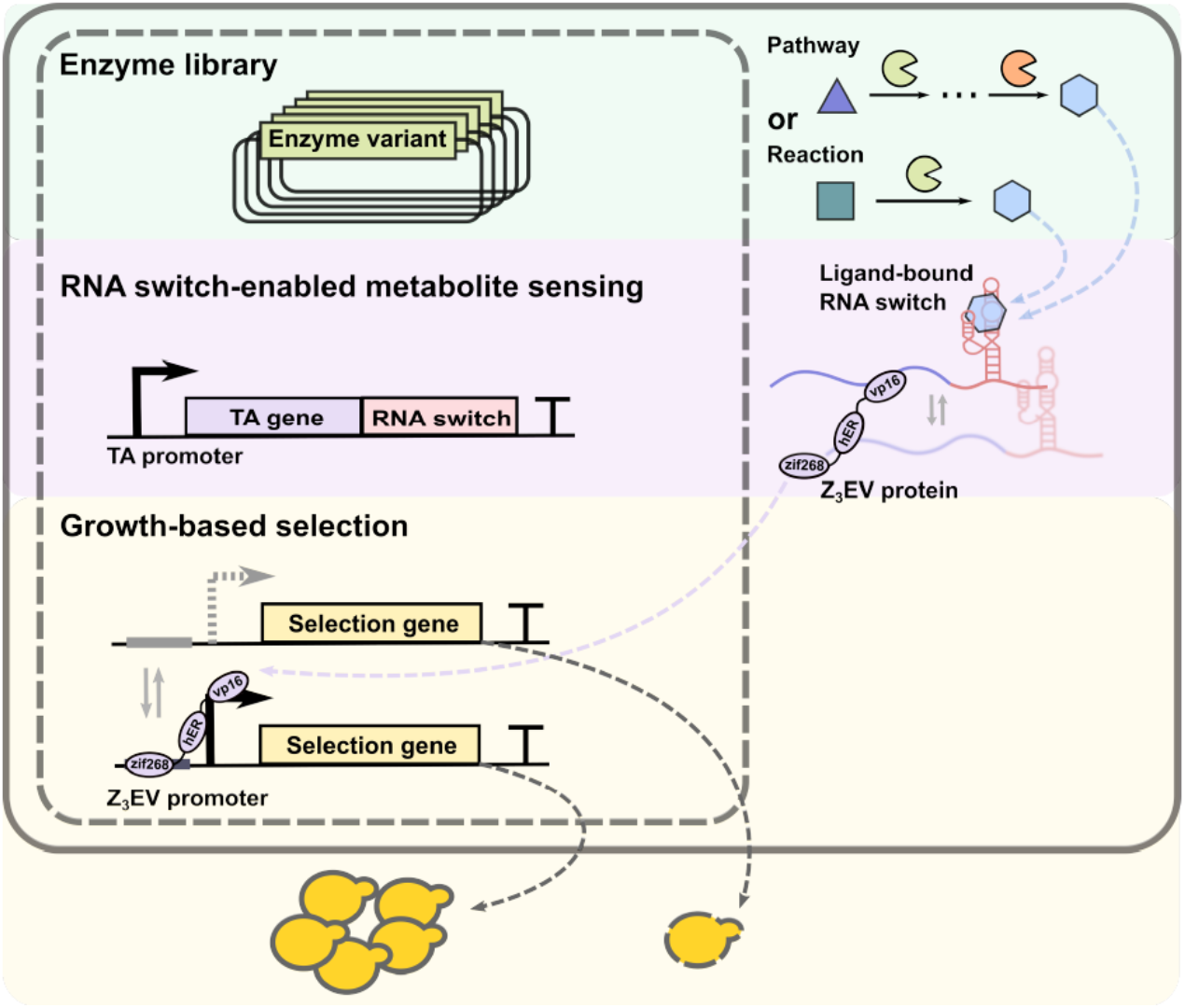
Schematic of the RNA switch-based selection system. Square-shaped solid line represents membrane of the cell, and square-shaped dotted line represents membrane of the nucleus. DNA information of an enzyme variant is encoded on a plasmid in the nucleus. DNA of the enzyme variant is transcribed and translated into the enzyme protein, which catalyzes a reaction leading to production of a key metabolite. The key metabolite is sensed by an RNA switch, which regulates the expression of a transactivator protein. The transactivator protein binds to the transactivated promoter, which regulates the expression of a selection gene. Expression of the selection gene determines cell fate.

## RESULTS

### Design and optimization of an RNA switch-based gene activation system

We sought to engineer a genetic selection system that would be able to convert changes in enzyme activity (or small molecule production) to differences in cell doubling time. As a first step, we designed a genetic system to increase the dynamic range of gene expression from a small molecule-responsive RNA switch, by adapting a previously described three-stage (sensing, processing, actuating) gene activation system^12^. The three-stage gene activation system uses a small molecule-responsive RNA switch to sense a small molecule input signal (sensing), i.e., intracellular concentration of a target metabolite; the RNA switch regulates the expression of a transcription factor protein (processing), which in turn controls expression of a reporter protein, indicated as an output signal, via a transcription factor-induced promoter (actuating). We designed a gene activation system based on this architecture using a recently described artificial transcription factor protein, Z_3_EV, and the corresponding Z_3_EV-inducible promoters (Z_3_EVp)^13^. The Z_3_EV promoter system is reported to be fast-acting (i.e., took less than 4 hours to achieve maximum reporter protein expression) and exhibits a low basal expression level in the absence of the inducer, both of which would provide advantages over the Tet-On promoter system that was adapted in the earlier design^12,14^.

We first characterized the activities of three Z_3_EV promoters, with three (Z_3_EV3p), five (Z_3_EV5p), and six (Z_3_EV6p) Z_3_EV protein binding sites, respectively (Figure 2A). We constructed low-copy plasmids encoding the Z_3_EV protein (pCS4580) and Z_3_EV3/5/6p-regulated yEGFP (pCS4581-4583). A wild-type yeast strain (CEN.PK2; MATα ura3-52, trp1-289; leu2-3/112, his3Δ1, MAL2-8C, SUC2) was co-transformed with the plasmid encoding the Z_3_EV protein (pCS4580) and the plasmid encoding either Z_3_EV3p (pCS4581), Z_3_EV5p (pCS4582), or Z_3_EV6p (pCS4583). The transformed strains were grown in synthetic dropout medium and induced with β-estradiol concentrations ranging from 0 to 1000 nM at an initial culture optical density at 600 nm (OD600) of 0.04. The median and standard deviation of fluorescence protein (yEGFP) expression level in the population of yeast cells was quantified using flow cytometry after 6 hours of growth in the presence of the inducer at 30°C. The maximum activation ratio was determined as a ratio of median yEGFP expression levels obtained at 1 and 0 mM β-estradiol concentrations. The Z_3_EV3p, Z_3_EV5p, and Z_3_EV6p systems exhibited maximum activation ratios of 5.08, 48.80, and 60.58, with basal EGFP expression levels of 14.28, 1.68 and 1.84, respectively (Figure 2B). Z_3_EV3p, Z_3_EV5p, and Z_3_EV6p each reached 51%, 48% and 62% activation at 10 nM β-estradiol, respectively. The observation is consistent with previous data where Z_3_EV6p shows the highest dynamic range^15^. We also confirmed that the system is fully activated at greater than 100 nM β-estradiol.

**Figure 2:**
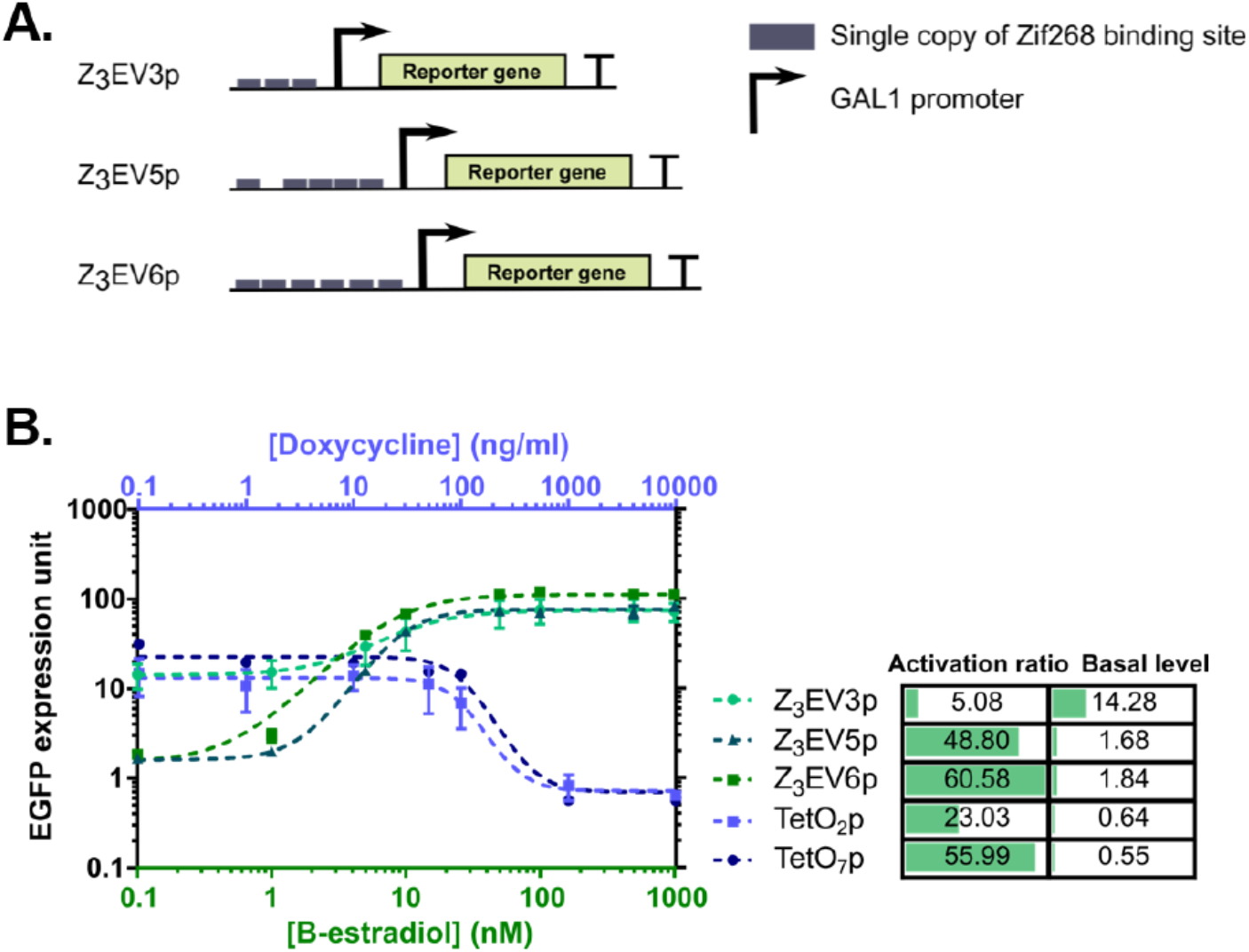
Construct schematic and activation ratios of Z_3_EV promoters. **A.** Construct designs for Z_3_EV promoters with three (Z_3_EV3p), five (Z_3_EV5p), or six (Z_3_EV6p) Z_3_EV protein binding sites. **B.** The activation ratios and basal expression levels of Z_3_EV promoters are compared to those of the well-characterized Tet-Off promoters.

In addition, we observed that the standard deviation in yEGFP expression for the three Z_3_EV promoters increases with β-estradiol concentration. For example, we observed 21.6%, 36.6%, and 82.7% increase in standard deviations in yEGFP expression for Z_3_EV3p, Z_3_EV5p, and Z_3_EV6p, respectively, between measurements in β-estradiol concentration at 1 nM and 5 nM, and 11.0%, 24.5%, and 26.3% increase in standard deviations between measurements in β-estradiol concentration at 10 nM and 50 nM (Figure 3A). The data indicates that higher levels of induction of the Z_3_EV promoters result in a greater level of noise (as measured by standard deviation in yEGFP expression of cell population originated from a single colony, treated by the same experimental condition).

**Figure 3:**
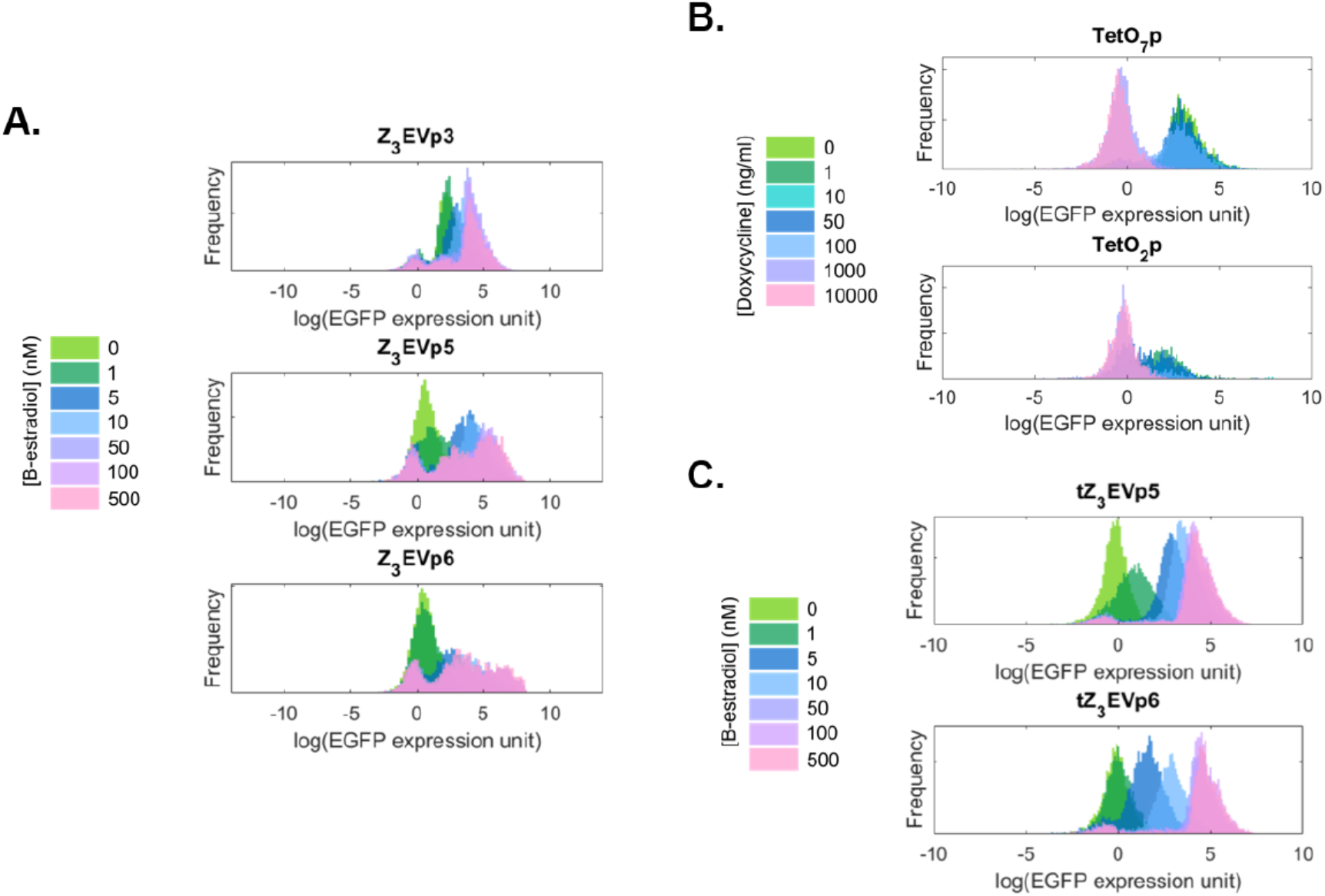
Noise profiles of Z_3_EV promoters, Tet-Off promoters and tZ_3_EV promoters. Noise profiles of **A.** Z_3_EV promoters induced by a range of β-estradiol concentrations, **B.** Tet-Off promoters induced by a range of doxycycline concentrations, and **C.** optimized tZ_3_EV promoters induced by the same range of β-estradiol concentrations as for Z_3_EV promoters. Noise profiles of each promoter are measured from yEGFP expression levels of a cell population originated from a single colony.

Since a high level of cell-to-cell variability in gene expression may compromise the phenotype to be displayed by the system, we attempted to optimize the Z_3_EV promoters to reduce their noise in gene expression. In a previously designed TetO promoter, an ADH1 terminator (ADH1t) sequence was placed upstream of the TetO promoter to avoid possible read-through from the plasmid sequences^16^. We observed that the TetO system incorporated with the terminator showed only 50% of the standard deviation in EGFP expression when fully expressed as compared to the fully induced Z_3_EV promoters Z_3_EV5p and Z_3_EV6p (Figure 3B). We hypothesized that the ADH1t sequence could potentially reduce gene expression noise by preventing read-through from sequences upstream of the promoter. We then attempted to demonstrate reduction in Z_3_EV promoter noise through integration of a terminator sequence upstream of the Z_3_EV protein binding sites.

We modified the constructs harboring Z_3_EV5p (pCS4582) and Z_3_EV6p (pCS4583) promoters by inserting the ADH1t sequence upstream of the Z_3_EV protein binding sites for each construct, resulting in the modified constructs tZ_3_EV5p (pCS4587) and tZ_3_EV6p (pCS4588), respectively. A wild-type yeast strain (CEN.PK2) was co-transformed with the plasmid encoding the Z_3_EV protein (pCS4580) and the plasmid encoding Z_3_EV5p (pCS4582), Z_3_EV6p (pCS4583), tZ_3_EV5p (pCS4587) or tZ_3_EV6p (pCS4588). The transformed strains were grown in synthetic dropout medium and induced with β-estradiol concentrations ranging from 0 to 500 nM at an initial culture OD600 culture of 0.04. The median and standard deviation in yEGFP expression in the population of yeast cells was quantified using flow cytometry after 6 hours of growth in the presence of the inducer at 30°C. The standard deviation in yEGFP expression in the sample cell population was consistently reduced by greater than 50% in tZ_3_EV5p compared to Z_3_EV5p and greater than 40% in tZ_3_EV6p compared to Z_3_EV6p when induced by β-estradiol of concentrations greater than 50 nM (Figure 3C), which confirmed that incorporating a terminator sequence upstream of a Z_3_EV promoter has effectively reduced the noise in its expression at high β-estradiol induction levels. In addition, the tZ_3_EV5p and tZ_3_EV6p systems exhibited maximum activation ratios of 62.67 and 67.81, with basal yEGFP expression levels of 0.98 and 0.44, respectively (Figure 4), which confirmed that the upstream terminator sequence did not compromise the dynamic range of the Z_3_EV promoters.

**Figure 4:**
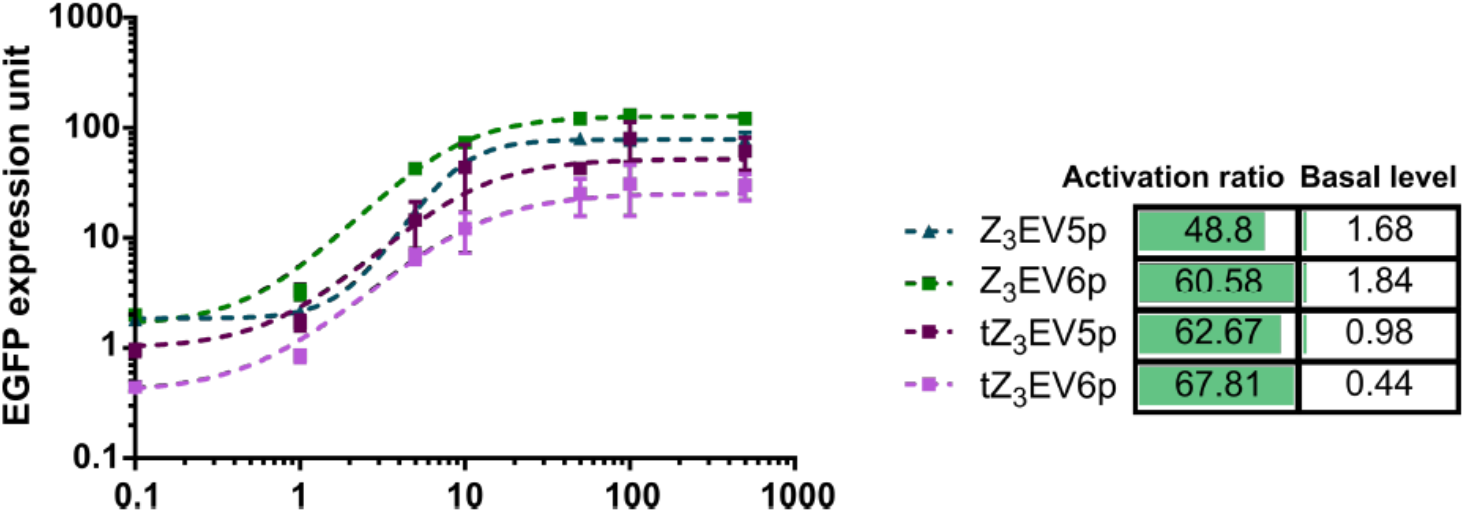
Dynamic range of optimized tZ_3_EV promoters, compared to that of Z_3_EV promoters.

We next characterized the performance of the Z_3_EV protein-based three-stage gene activation system, under the regulation of the optimized tZ_3_EV protein and a theophylline-responsive RNA switch. We first tested the system response with fed theophylline (Figure 5). We designed three constructs for reporting yEGFP signal: (i) a construct without the transcription factor processing stage, harboring the RNA switch at the 3’ untranslated region (UTR) of the yEGFP reporter gene; (ii) a construct with the transcription factor expression, harboring the RNA switch at the 3’ UTR of both the transactivator gene and the reporter gene; (iii) a construct with the transcription factor expression, harboring the RNA switch only at the 3’ UTR of the transactivator gene (Figure 6A).

**Figure 5:**
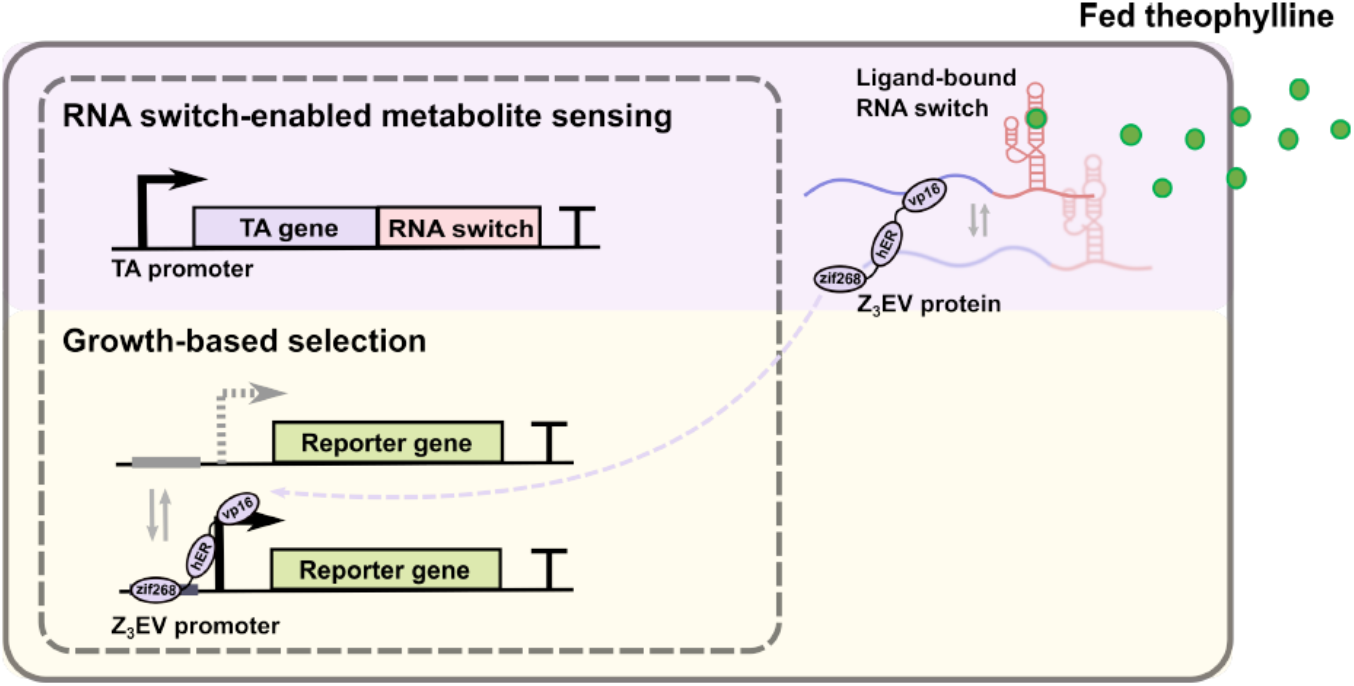
System schematic for characterization of Z_3_EV protein and RNA switch-based regulatory constructs. Theophylline is fed into the system to titrate yEGFP reporter gene expression via a theophylline-responsive RNA switch. The theophylline-responsive RNA switch regulates the expression of the Z_3_EV protein which activates the Z_3_EV promoter including yEGFP reporter gene expression.

**Figure 6:**
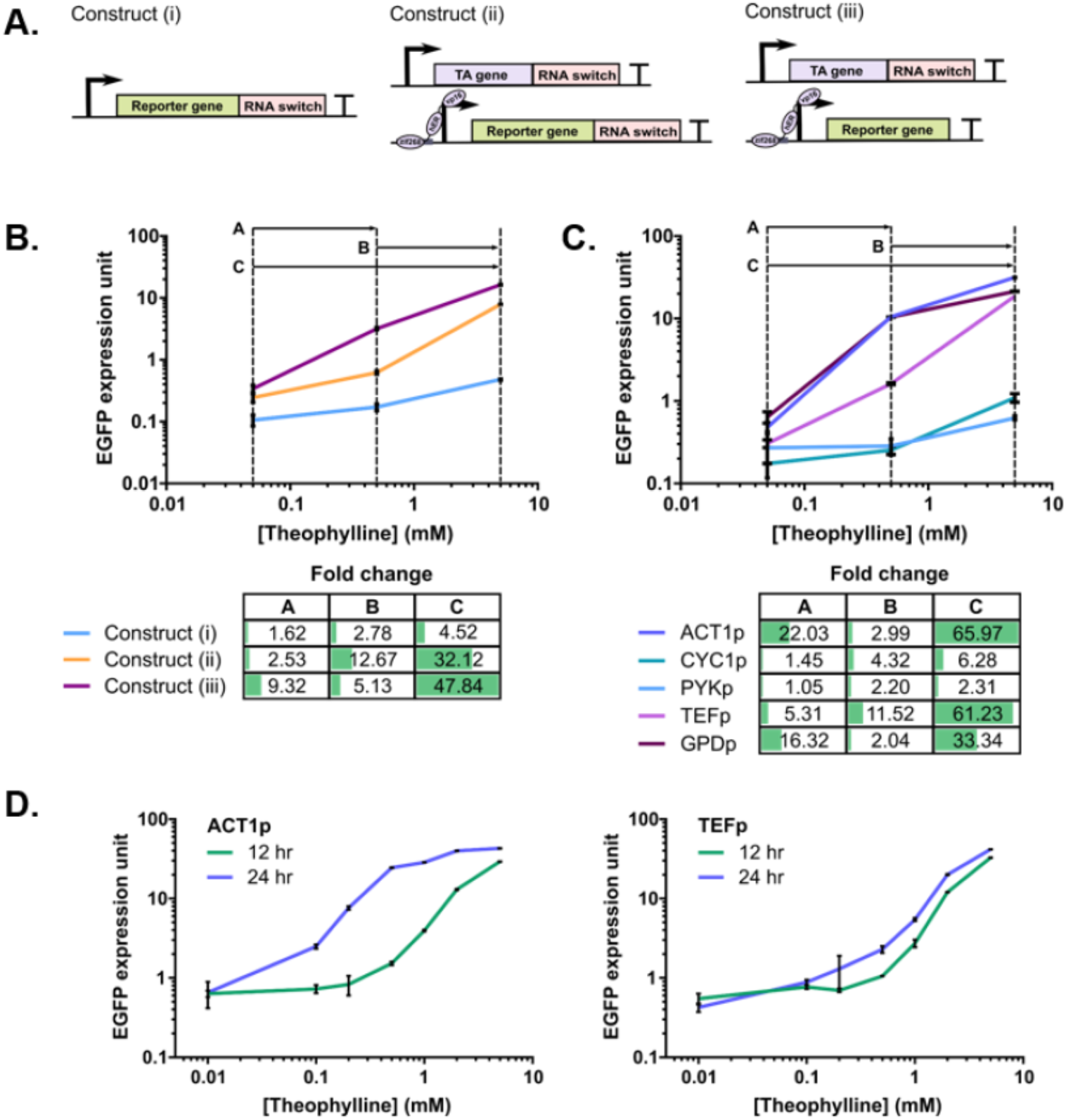
Design of the RNA switch-based expression platform. **A.** Gene expression constructs for RNA switch-based regulation with or without incorporation of a transactivator (TA). **B.** Comparison in the dynamic range of the gene expression constructs. **C.** Comparison in dynamic range of construct (iii) using different promoters for TA gene expression. **D.** Characterization of the temporal change in dynamic range construct (iii) using ACT1p or TEFp for TA gene expression.

We cloned three additional constructs on low-copy plasmids: an ACT1p-regulated yEGFP expression cassette with RNA switch at 3’ UTR (pCS4589), an ACT1p-regulated Z_3_EV protein expression cassette with RNA switch at 3’ UTR (pCS4590), and a tZ_3_EV6p-regulated yEGFP expression cassette with RNA switch at 3’ UTR (pCS4591). We integrated the linear DNA template of the expression cassette PCR-amplified from pCS4589 into CEN.PK2, resulting in CSY1362 (i). Using a similar technique, we integrated the expression cassette from pCS4590 into CEN.PK2, resulting in CSY1363. We next integrated the expression cassette on pCS4591 into CSY1363, resulting in CSY1364 (ii), and the expression cassette on pCS4988 (tZ_3_EV6p-regulated EGFP expression without RNA switch at 3’ UTR) into CSY1363, resulting in CSY1365 (iii). The strains were grown in synthetic complete medium with 0, 0.5, or 5 mM theophylline and induced with 0.5 μM β-estradiol at an initial culture OD600 of 0.04. The median yEGFP expression level in cell population was quantified using flow cytometry after 12 hours of growth at 30°C. Constructs (i)-(iii) exhibited dynamic ranges of 1.62, 2.53, and 9.32, respectively, when induced by 0.5 mM (versus 0 mM) theophylline, and dynamic ranges of 4.52, 32.1, and 47.8, respectively, when induced by 5 mM (versus 0 mM) theophylline (Figure 6B). Construct (iii) shows 5.75-fold and 10.6-fold higher dynamic range compared to construct (i) when induced by 0.5 mM and 5 mM theophylline, respectively, which indicates that the Z_3_EV protein-based gene activation system has effectively enhanced the dynamic range of the RNA switch compared to a design without the transcription factor (i). In addition, construct (iii) shows 3.68-fold and 1.48-fold higher dynamic range compared to construct (ii) when induced by 0.5 mM and 5 mM theophylline, respectively, which indicates that the Z_3_EV protein-based gene activation system exhibited a greater dynamic range when harboring the RNA switch only at 3’ UTR of the transactivator gene (iii) (Figure 6B).

Finally, we examined the effects of the transactivator promoter strength on the dynamic range observed with the Z_3_EV protein and RNA switch-based gene activation system. The tZ_3_EV6p-regulated EGFP expression cassette was integrated into the wild-type yeast strain CEN.PK2, resulting in CSY1366. We designed the Z_3_EV protein expression cassette using construct (iii) (RNA switch only at the 3’ UTR of the transactivator gene), regulated by promoters of varying strengths (low to high): CYC1p, PYKp, TEFp, GPDp. We PCR amplified the promoter sequences from pCS2659 (CYC1p), pCS2664 (PYKp), pCS2657 (TEFp), and pCS2656 (GPDp), and Z_3_EV protein sequence with RNA switch at 3’ UTR from pCS4590. We integrated the overlap PCR fragments of complete Z_3_EV protein expression cassettes into yeast strain CSY1366 (strain harboring tZ_3_EV6p-regulated EGFP expression cassette), resulting in CSY1367 (CYC1p), CSY1368 (PYKp), CSY1369 (TEFp), and CSY1370 (GPDp). The strains plus the strain incorporating ACT1p (CSY1365) were grown in synthetic complete medium with 0, 0.5, or 5 mM fed theophylline and induced with 0.5 μM β-estradiol at an initial culture OD600 of 0.04. The yEGFP expression level in the population of yeast cells was quantified using flow cytometry after 12 and 24 hours of growth at 30°C. Medium to high activity promoters ACT1p and TEFp-regulated constructs exhibited the highest and similar overall dynamic ranges (53.6 and 51.3, respectively, at 12 hour) when induced by 5 mM theophylline, although ACT1p-regulated construct showed slightly higher dynamic range (6.36) compared to that of TEFp-regulated construct (3.08) when induced by 0.5 mM theophylline at 12 hour (Figure 6C). TEFp results in the highest dynamic range between 0.5 mM and 5 mM theophylline concentrations (16.6), and greater stability in expression levels over time, compared to ACT1p, for which the basal expression level increases over time (Figure 6D). Since our goal is to select for high enzyme activities, the dynamic range between 0.5 mM and 5 mM theophylline titrations was a determining factor to select the TEFp-regulated construct for further characterizations of the growth regulatory genes in the selection system.

### Characterization of auxotrophic genes for yeast growth regulation

The use of a transcription factor for enhancing RNA switch-based regulation of gene expression provides a starting point for the design of a selection system that links enzyme activity to yeast growth. We next replaced the yEGFP reporter gene in the transcription factor-based RNA switch activation system with a selection gene and tested the resulting strains for growth regulation (Figure 7). Here, we first tested a group of auxotrophic genes, i.e., *URA3, SpHIS5, LEU2*, and *TRP1*, for yeast growth regulation in synthetic amino acid dropout medium lacking either uracil, histidine, leucine, or tryptophan. Previous study reported evolved variants of a caffeine demethylase (CDM) enzyme, which exhibited different levels of activity converting caffeine to theophylline^9^. We used these enzyme variants to test the ability of each selection gene to differentiate yeast growth using the Z_3_EV protein and RNA switch-based gene activation system, e.g., a high-producing enzyme variant is expected to result in a higher growth rate compared to a low-producing variant.

**Figure 7:**
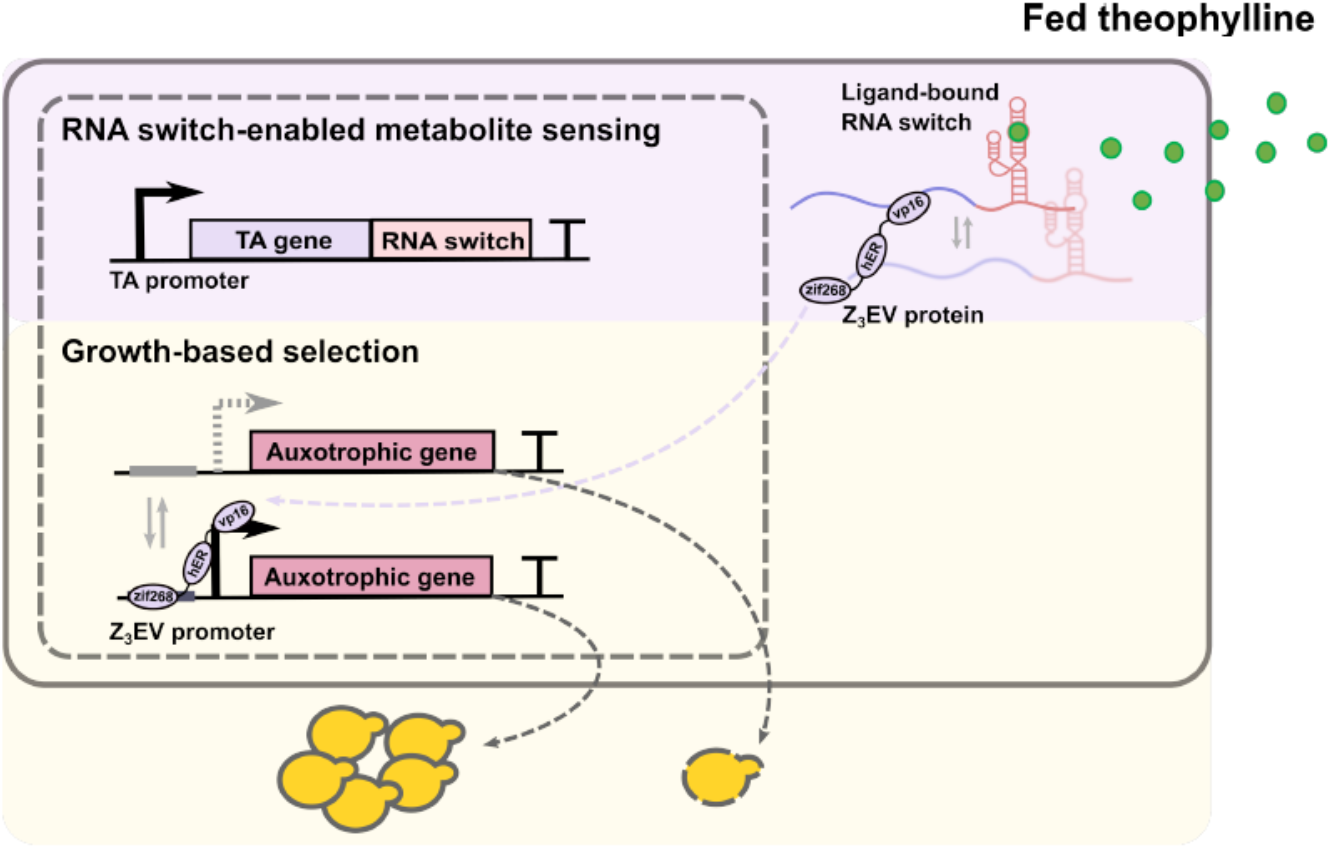
System schematic for characterization of auxotrophic gene activity with a ligand-responsive RNA switch and fed ligand. Theophylline is fed into the system to titrate auxotrophic gene expression via a theophylline-responsive RNA switch. The theophylline-responsive RNA switch regulates the expression of the Z_3_EV protein which activates the Z_3_EV promoter inducing auxotrophic gene expression.

To facilitate the characterization of selection genes in our system, we examined the correlation between the CDM variant activities in the presence of 1 mM fed caffeine and various fed theophylline concentrations. Low-copy plasmids encoding expression constructs for the CDM variants exhibiting low (CDM1/2), medium (CDM4/5), or high (CDM7/8) activities and a negative control (inactive *ccdB*) were transformed into CSY1365 (reporter strain with Z_3_EV protein expression regulated by RNA switch at 3’ UTR and tZ_3_EV6p-regulated yEGFP expression). The resulting strains were grown in synthetic dropout medium with either 1 mM caffeine or 0, 0.5, or 5 mM theophylline and induced with 0.5 μM β-estradiol at an initial culture OD600 of 0.04. The yEGFP median expression level in the population of yeast cells was quantified using flow cytometry after 24 hours of growth at 30°C. From the yEGFP expression, we categorized the enzyme variants CDM1/2/4/5 as low-producers (which correspond to EGFP levels between that from cells grown in 0 to 0.5 mM fed theophylline) and CDM7/8 as high-producers (which correspond to yEGFP levels higher than those exhibited from cells grown in 0.5 mM fed theophylline) (Figure 8A). To further confirm activities of the CDM variants, theophylline production by CDM5/7/8 was also quantified in the growth medium by LC-MS analysis (Figure 8B). By correlating CDM enzyme variant activities from fed theophylline concentrations, we can test the system, i.e. measure yeast growth, by titrating selection genes with an expected level of CDM activity (low or high) by feeding different concentrations of theophylline, without the need to transform each CDM variant into each selection strain.

**Figure 8:**
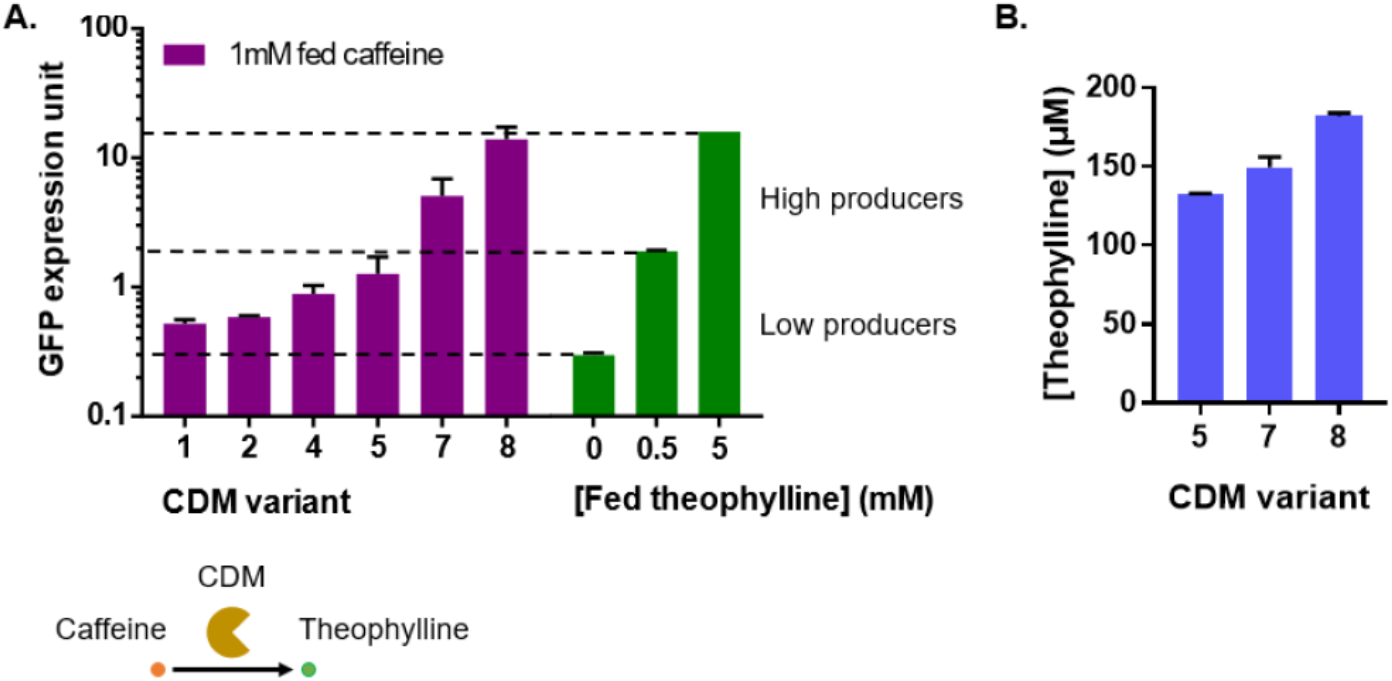
Characterization of CDM variant activities. **A.** yEGFP expression resulting from intracellular theophylline production by CDM variant is compared to yEGFP expression resulting from theophylline fed into yeast medium. A CDM variant catalyzes the conversion of caffeine to theophylline. Caffeine is fed into yeast medium for characterization of CDM activity. **B.** Characterization of theophylline production in yeast medium by CDM 5/7/8 via LC-MS.

We first characterized the auxotrophic genes *URA3, SpHIS5, LEU2*, and *TRP1* and titrated the system to measure yeast growth at different fed theophylline levels (0, 0.5, and 5 mM). We created a base strain harboring TEFp-regulated Z_3_EV protein expression cassette with RNA switch at 3’ UTR by PCR amplifying the expression cassette from CSY1369, and integrating the linear DNA template into a wild-type strain (CEN.PK2), resulting in CSY1371. We next integrated tZ_3_EV6p-regulated expression cassettes of the auxotrophic genes into CSY1371, resulting in selection strains encoding *URA3* (CSY1372), *SpHIS5* (CSY1373), *LEU2* (CSY1374), and *TRP1* (CSY1375). The resulting strains were grown in synthetic dropout medium with 0, 0.5, or 5 mM theophylline and induced with 0.5 μM β-estradiol at an initial culture OD600 of 0.04. The OD600 of each strain was measured at 0, 6, 12, and 24 hours using a plate reader over the course of the assay. Strains harboring the auxotrophic genes *URA3, SpHIS5*, and *LEU2* all exhibited titratable growth by theophylline, resulting in 8.05, 30.5, and 17.4-fold higher OD600 with 5 mM fed theophylline compared to 0 mM theophylline after 24 hours (Figure 9A). Furthermore, *SpHIS5* and *LEU2* exhibited low growth when fed with 0.5 mM theophylline, and the differences in OD600 between 0.5 and 0 mM fed theophylline levels are negligible. The results indicate that a single CDM variant achieving production levels comparable to 5 mM fed theophylline can potentially dominate the cell population by 70%, enriched from an initial population of 10,000 low activity CDM variants (production level comparable to 0.5 mM fed theophylline) using the *SpHIS5* selection strain in 72 hours, without considering nutrient depletion in medium or culture reaching saturation. The auxotrophic gene *TRP1* showed significant growth in the absence of theophylline. This growth is likely resulted from a basal expression of the Z_3_EV promoter, due to which uninhibited growth was measured in the absence of the inducer in dropout medium (Figure 9B). The results indicate that *SpHIS5* in the context of the described selection system resulted in low basal growth and high fold change in culture OD600 when induced by theophylline.

**Figure 9:**
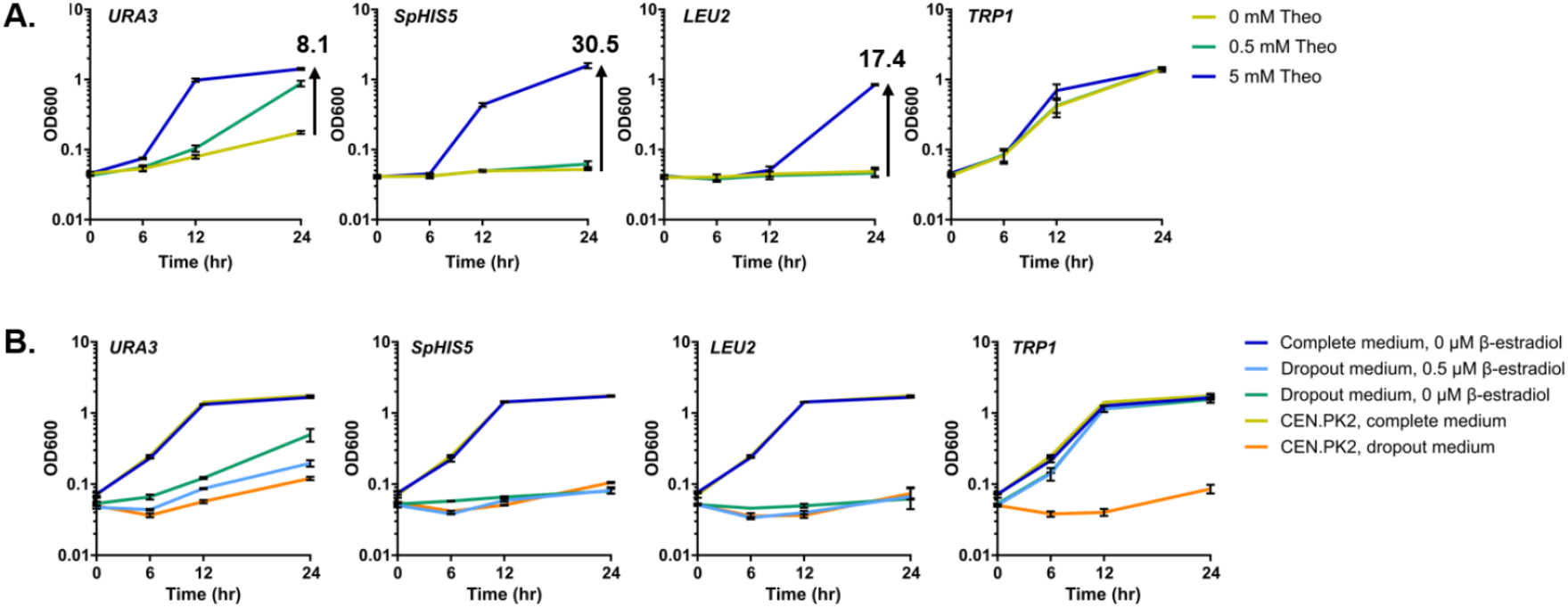
Growth regulation by auxotrophic genes using a theophylline responsive RNA switch. **A.** Growth of selection strains expressing either one of the four auxotrophic genes (*URA3, SpHIS5, LEU2*, or *TRP1*) is measured by OD600 at four time points, and induced by 0, 0.5, or 5 mM theophylline. **B.** Basal growth resulted from the Z_3_EV protein and RNA switch-based expression system is measured by growth of each selection strain in the absence of the inducer (β-estradiol) in synthetic dropout medium. CEN.PK2 is used as an auxotrophic gene negative control.

To validate the use of an auxotrophic gene for enriching high-producing enzyme variants from a cell population, we used *SpHIS5* gene to demonstrate the enrichment of a high-producing variant of CDM (CDM8) among a random mutagenesis library of CDM5. An enzyme variant is transformed into the yeast cell, which produces theophylline from caffeine fed into yeast medium (Figure 10). Intracellular theophylline production levels are then detected by an RNA switch, which titrates the expression of the auxotrophic gene via induction by the transactivator protein.

**Figure 10:**
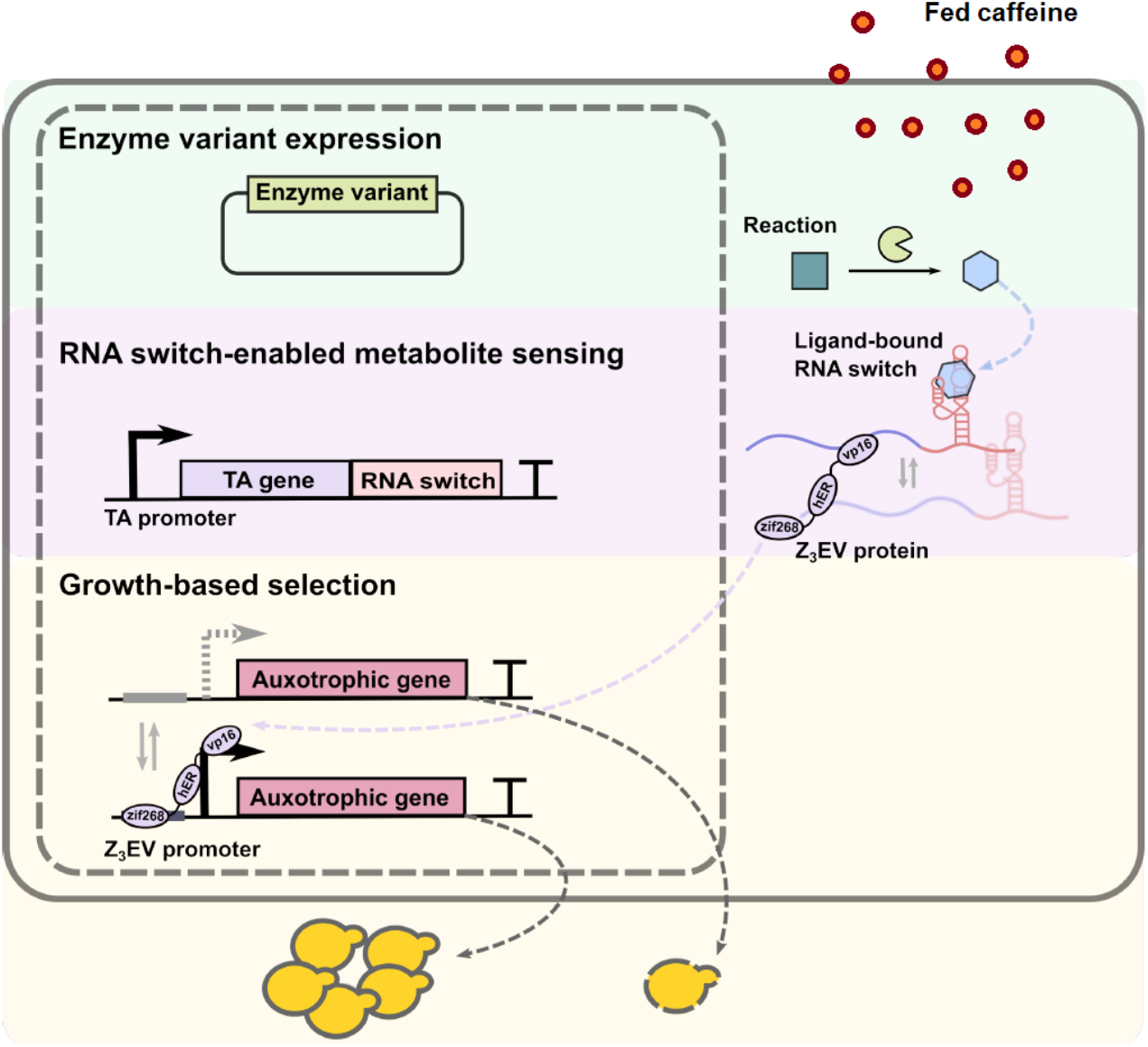
System schematic for characterization of auxotrophic gene activity with a metabolitesensing RNA switch and an enzyme variant producing the metabolite. Theophylline is converted by CDM from caffeine fed into yeast medium. The theophylline-responsive RNA switch regulates the expression of Z_3_EV protein which activates the Z_3_EV promoter inducing auxotrophic gene expression.

We first PCR-amplified the targeted genetic sequence of CDM5 enzyme variant from pCS2165 using GeneMorph II kit, and generated a library of PCR fragments harboring 3-5 mutations per gene. We next digested the low-copy expression vector encoded by pCS2168 using BlpI and EcoRI, and gel purified the digested vector products. We transformed library PCR product and digested vector product into the competent cells of CSY1373 by following an electroporation protocol and seeded transformed cells in 50 ml uracil dropout medium for 16 hours of recovery at 30°C. In addition, we inoculated an overnight culture of the CDM8 enzyme variant in uracil dropout medium and incubated at 30°C for 16 hours. We measured OD600 of both the recovered CDM5 library culture and the CDM8 variant culture. We spined down 10 ml of CDM5 library culture, resuspended the culture in 1 ml of uracil dropout medium, and spiked in 0.1% of CDM8 enzyme variant overnight culture based on OD600 ratio between CDM8 culture and unconcentrated CDM5 library culture. We inoculated the 1 ml of culture mix into 50 ml of fresh uracil and histidine dropout medium in a 250 ml flask, induced by 0.5 μM β-estradiol. We incubated the flask culture at 30°C overnight and 50-fold back diluted the culture into 50 ml of fresh medium after 24 hours of growth (1 ml overnight culture in 50 ml fresh medium). The back dilution is repeated for another 5 rounds with a dilution rate of 100-fold (500 μl overnight culture in 50 ml fresh medium), and we sampled culture at the start of round 1 (denoted at ‘round 0’), and at the end of round 6. CDM8 variant harbors three additional mutations compared to the CDM5 variant, located at base positions 332, 593, and 1377 in the CDM gene sequence. Our data shows that the mutations in CDM8 are enriched to around 20% of the entire population after six rounds of back dilution (Figure 11).

**Figure 11:**
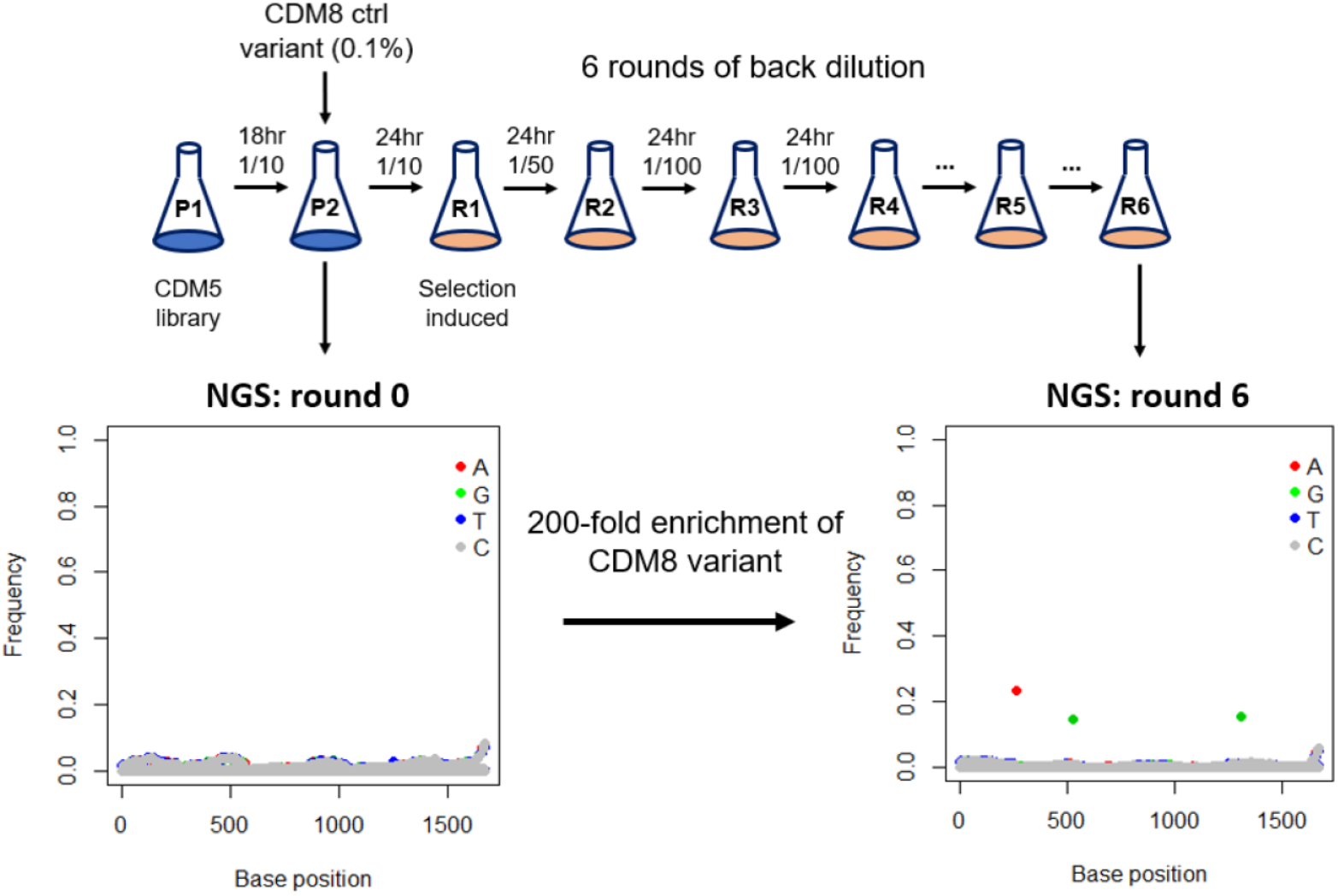
CDM8 enrichment using the auxotrophic gene *SpHIS5*. A high-producing CDM variant, CDM8, is spiked into a mutagenesis library of low-producing CDM5 variants with an initial presence of 0.1% of the entire population. The high-producing variant CDM8 is enriched to around 20% of the entire population after iterations of growth in flask culture.

In this enrichment experiment, only the high-producing variant CDM8 was successfully enriched, and none of the CDM5 variants from the random mutagenesis library have been enriched. Two potential explanations could be given for the lack of distinguished growth from low-producing variant library: (i) the current selection pressure by auxotrophy alone is too strong and therefore has been fully inhibiting any variant growth; (ii) the selection pressure by auxotrophy alone enables equally low growth of the variants from the library and is not sufficient to differentiate small differences in variant activities. From the characterization data in Figure 9A, low growth was observed in individual colonies of the selection strain fed with 0.5 mM theophylline, which indicates a possibility of explanation (ii).

### Incorporation of an antibiotics-based selective pressure enables selection of low-producing variants

To further distinguish growth of low-producing enzyme variants, we examined the possibility of incorporating a second selective pressure, i.e., antibiotics, into the selection system. We first tested individual activities of four antibiotic resistance genes (*KanR, HygroR, NrsR*, and *CyhR*). In this system, the Z_3_EV protein-based transcription factor induces the expression of an antibiotic resistance gene under the regulation of tZ_3_EV6p promoter (Figure 12). Antibiotic is added to yeast medium, and the antibiotic resistance gene is titrated by the theophylline-responsive RNA switch with fed theophylline to monitor yeast cell growth. The antibiotic selection cassettes under the regulation of tZ_3_EV6p were chromosomally integrated into CSY1371 (TEF1p-regulated Z_3_EV protein expression cassette with RNA switch at 3’ UTR), resulting in selection strains encoding antibiotic resistance genes *KanR* (CSY1376), *HygroR* (CSY1377), *NrsR* (CSY1378) and *CyhR* (CSY1379). The resulting strains were grown in synthetic complete medium with 0.25 or 0.5 mM theophylline and induced with 0.5 μM β-estradiol at an initial culture OD600 of 0.04 and supplemented with the appropriate antibiotic, i.e., geneticin (CSY1376), hygromycin B (CSY1377), nourseothricin (CSY1378), or cycloheximide (CSY1379). For each antibiotic, CEN.PK2 was used as a negative control (absence of antibiotic resistance gene expression), induced and incubated under the same conditions. The OD600 was measured at 0, 6, 12, and 24 hours for CSY1376-1377 and at 0, 12, 24, 56 hours for CSY1378-1379 using a plate reader over the time course of the assay. *KanR* showed uninhibited growth with basal expression of tZ_3_EV6p in the absence of theophylline and was ineffective at distinguishing growth at different concentrations of theophylline, at either 100 or 200 μg/ml geneticin. The strain harboring the *HygroR* expression cassette exhibited a three-fold difference in OD600 at 24 hour when fed with 0.25 and 0.5 mM theophylline, supplied with 500 μg/ml hygromycin B. The strain harboring the *NrsR* expression cassette exhibited a 10-fold difference in OD600 at 56 hour when fed with 0.25 and 0.5 mM theophylline, supplied with 50 μg/ml nourseothricin. However, no growth was observed from the strain harboring the *CyhR* expression cassette at the concentrations of cycloheximide tested (Figure 13). The data indicates that both antibiotic resistance genes *HygroR* and *NrsR* show differences in growth with 0.25 and 0.5 mM fed theophylline levels mimicking low producing CDM variants; therefore, hygromycin B and are nourseothricin are chosen as the second selection pressure for distinguishing low-producers, in addition to auxotrophic selection pressure.

**Figure 12:**
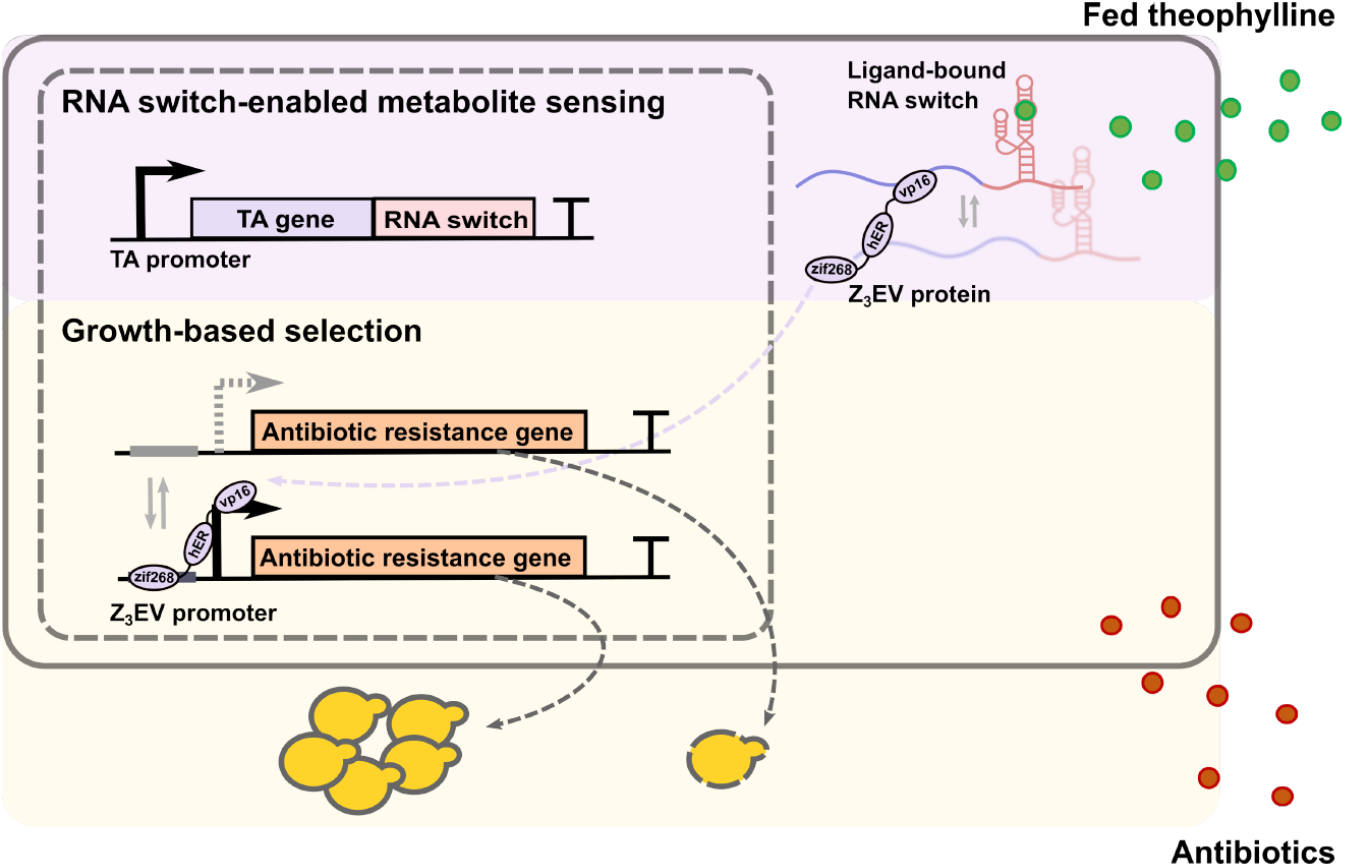
System schematic for characterization of antibiotic resistance gene activity with a theophylline-responsive RNA switch. Antibiotic is added to yeast medium. Theophylline is fed into the system to titrate antibiotic resistance gene expression via a theophylline-responsive RNA switch. The theophylline-responsive RNA switch regulates the expression of the Z_3_EV protein which activates the Z_3_EV promoter inducing antibiotic resistance gene expression.

**Figure 13:**
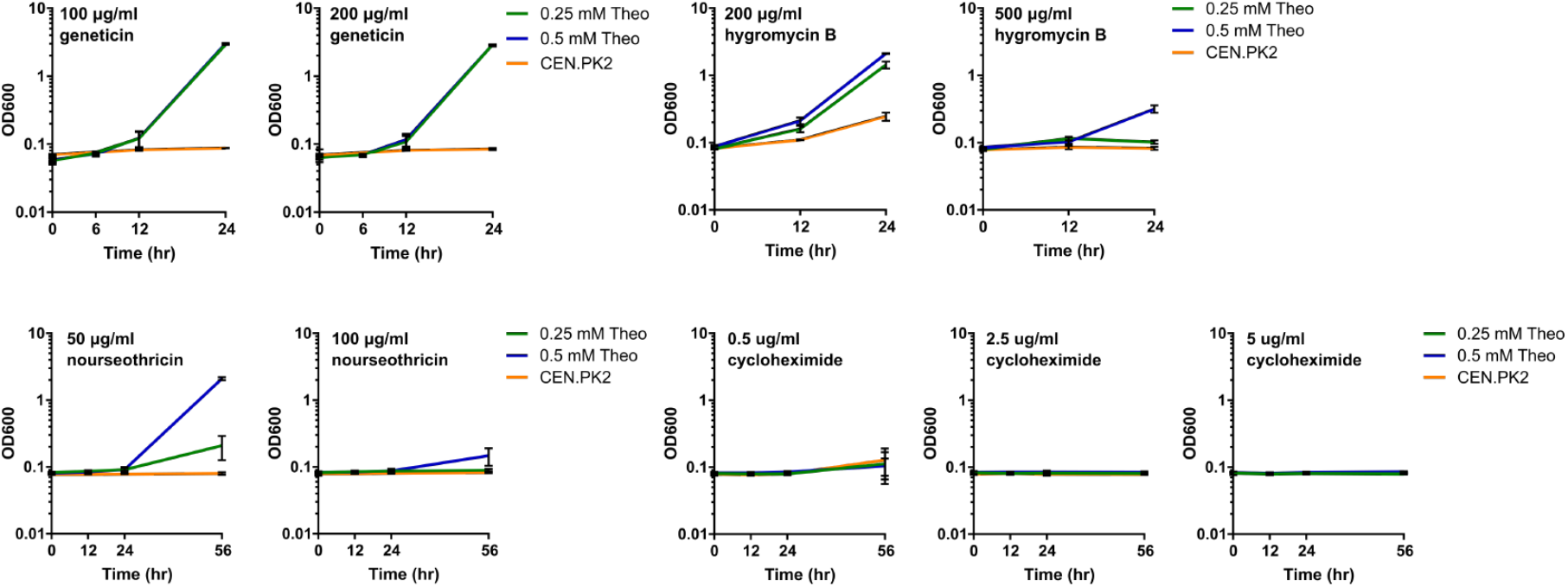
Growth regulation by antibiotic resistance genes using a theophylline-responsive RNA switch. Growth of selection strains expressing either one of the four antibiotic resistance genes (*KanR, HygroR, NrsR, CyhR*) is measured by OD600 at three or four time points, and induced by 0.25 or 0.5 mM theophylline. Corresponding antibiotic is added to yeast medium of each selection strain: geneticin for *KanR*.

We next characterized selection activities of both an auxotrophic gene and an antibiotic resistance gene when regulated by the Z_3_EV protein and RNA switch-based gene activation system. Here, we termed the selection system incorporating both an auxotrophic and an antibiotic resistance gene a dual selection system. Instead of using the Z_3_EV protein-based transcription factor to induce the expression of one selection gene, we incorporated a second selection gene expression cassette under the regulation of tZ_3_EV6p promoter (Figure 14). As such, both the auxotrophic gene and the antibiotic resistance gene are titrated by the theophylline-responsive RNA switch with fed theophylline to monitor yeast growth.

**Figure 14:**
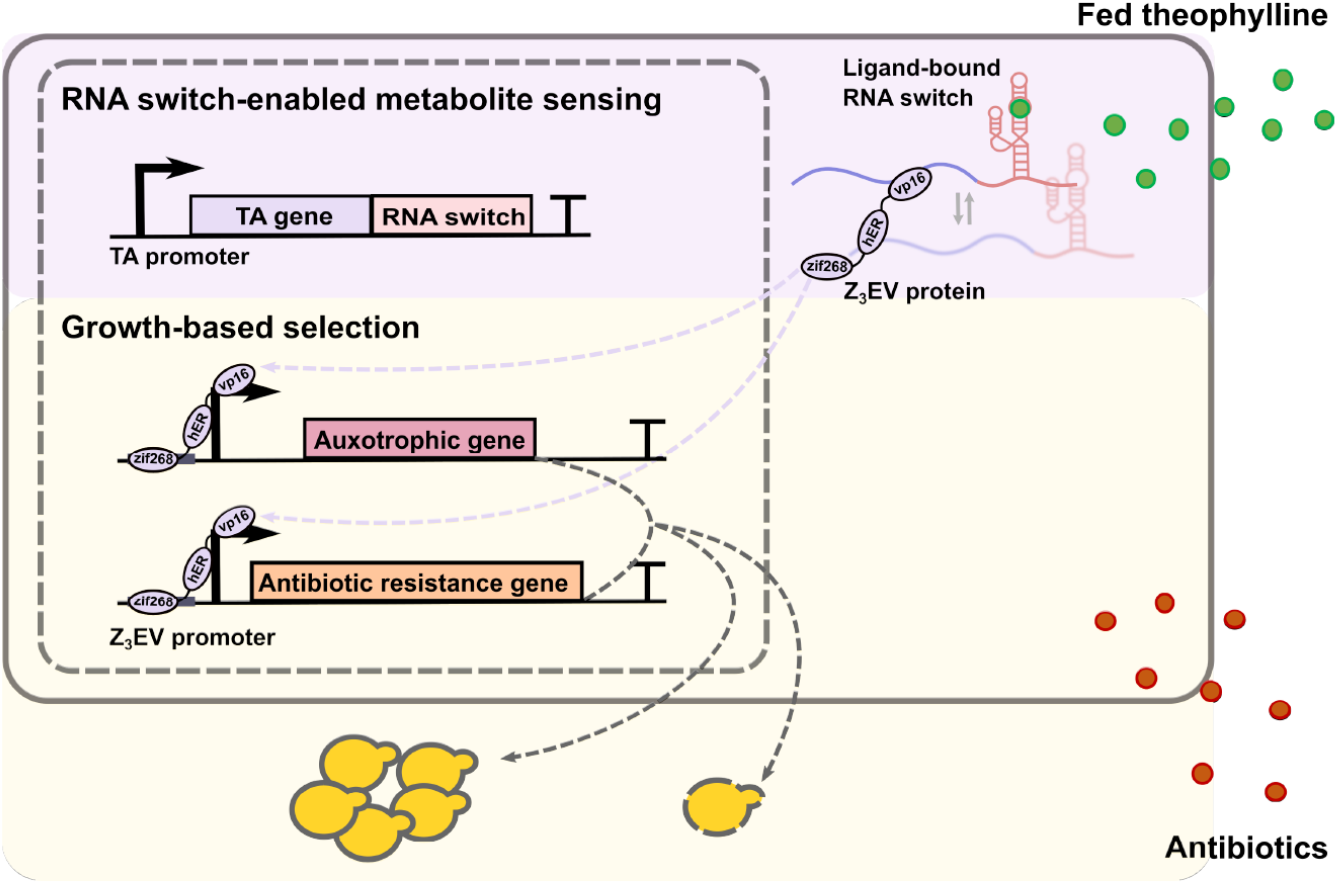
System schematic for incorporating a second selection pressure introduced by antibiotics. Antibiotic is added to yeast medium as a second selection pressure. Theophylline is fed into the system to titrate auxotrophic and antibiotic resistance gene expression via a theophylline-responsive RNA switch. The theophylline-responsive RNA switch regulates the expression of the Z_3_EV protein-based transcription factor which activates the Z_3_EV promoter inducing expression of both an auxotrophic gene and an antibiotic resistance gene.

We measured selectivity of the dual selection system by measuring its ability to eliminate growth of mid-to-low producing variants, as represented by 0.25 mM fed theophylline level, and to allow growth of mid-to-high producing variants, as represented by 0.5 mM fed theophylline level. We designed dual selection constructs based on pairing the expression of either one of the auxotrophic genes *SpHIS5, ScHIS3, LEU2*, or *URA3* with either *NrsR* or *HygroR* antibiotic resistance genes. Here, we tested an additional auxotrophic gene *ScHIS3*, which is a variant of imidazoleglycerol-phosphate dehydratase native to *S. cerevisiae*, and it is expected to catalyze the same reaction as *SpHIS5* from *Schizosaccharomyces pombe* in completing histidine biosynthesis in an auxotrophic strain, but could be at a higher rate. The two expression cassettes were iteratively integrated into CSY1371 (TEFp-regulated Z_3_EV protein expression cassette with RNA switch at 3’ UTR), resulting in the dual selection strains *SpHIS5/NrsR* (CSY1380), *SpHIS5/HygroR* (CSY1381), *ScHIS3/NrsR* (CSY1382), *ScHIS3/HygroR* (CSY1383), *ScHIS3/HygroR* (CSY1384), *LEU2/NrsR* (CSY1385), *LEU2/NrsR* (CSY1386), *LEU2/HygroR* (CSY1387), *URA3/NrsR* (CSY1388), and *URA3/HygroR* (CSY1389). The resulting strains were grown in synthetic dropout medium with 0, 0.25, or 0.5 mM theophylline and induced with 0.5 μM β-estradiol at an initial culture OD600 of 0.04 and supplemented with either 50 μg/ml nourseothricin or 500 μg/ml hygromycin B. The OD600 of each strain was measured at 0, 6, 12, 24, 30, and 48 hours using a plate reader over the time course of the assay. Yeast growth rate for each strain and condition was calculated based on the measured OD600 data.

Based on previously built correlation between CDM enzyme variant activities and fed theophylline concentrations, 0.5 mM theophylline represents a mid-to-high expected level of CDM activity (e.g., CDM6/7) and 0.25 mM theophylline represents a mid-to-low level of CDM enzyme activity (e.g., CDM2/4). We defined the differences in growth rates between 0.25 and 0.5 mM fed theophylline concentrations as the distinction factor of a selection strain, and the higher the distinction factor the better a selection is at inhibiting median level enzyme activities and promoting high level enzyme activities at the same time. Each of the single selection strain, *SpHIS5* (CSY1373), *ScHIS3* (CSY1390), *LEU2* (CSY1375), and *URA3* (CSY1372), resulted in a distinction factor of 2.52, 1.20, 4.23, and 2.84, respectively (Figure 15). The dual selection strains *SpHIS5/NrsR* (CSY1380) and *SpHIS5/HygroR* (CSY1381) exhibited distinction factors of 418 and 186, representing 70 to 160-fold improvement from the single selection strain *SpHIS5* (CSY1373). The dual selection strains *LEU2/NrsR* (CSY1386) and *SpHIS3/HygroR* (CSY1381) exhibited distinction factors of 15.3 and 6.53 respectively, representing 1.5 to 3.6-fold improvement from the single selection strain *LEU2* (CSY1375). No greater than two-fold improvement was observed in dual selection strains harboring *ScHIS3* or *URA3*. The results indicate that the dual selection strains harboring *SpHIS5* are most effective for distinguishing high producers (represented by 0.5 mM fed theophylline) from medium-low producers (represented by 0.25 mM fed theophylline).

**Figure 15:**
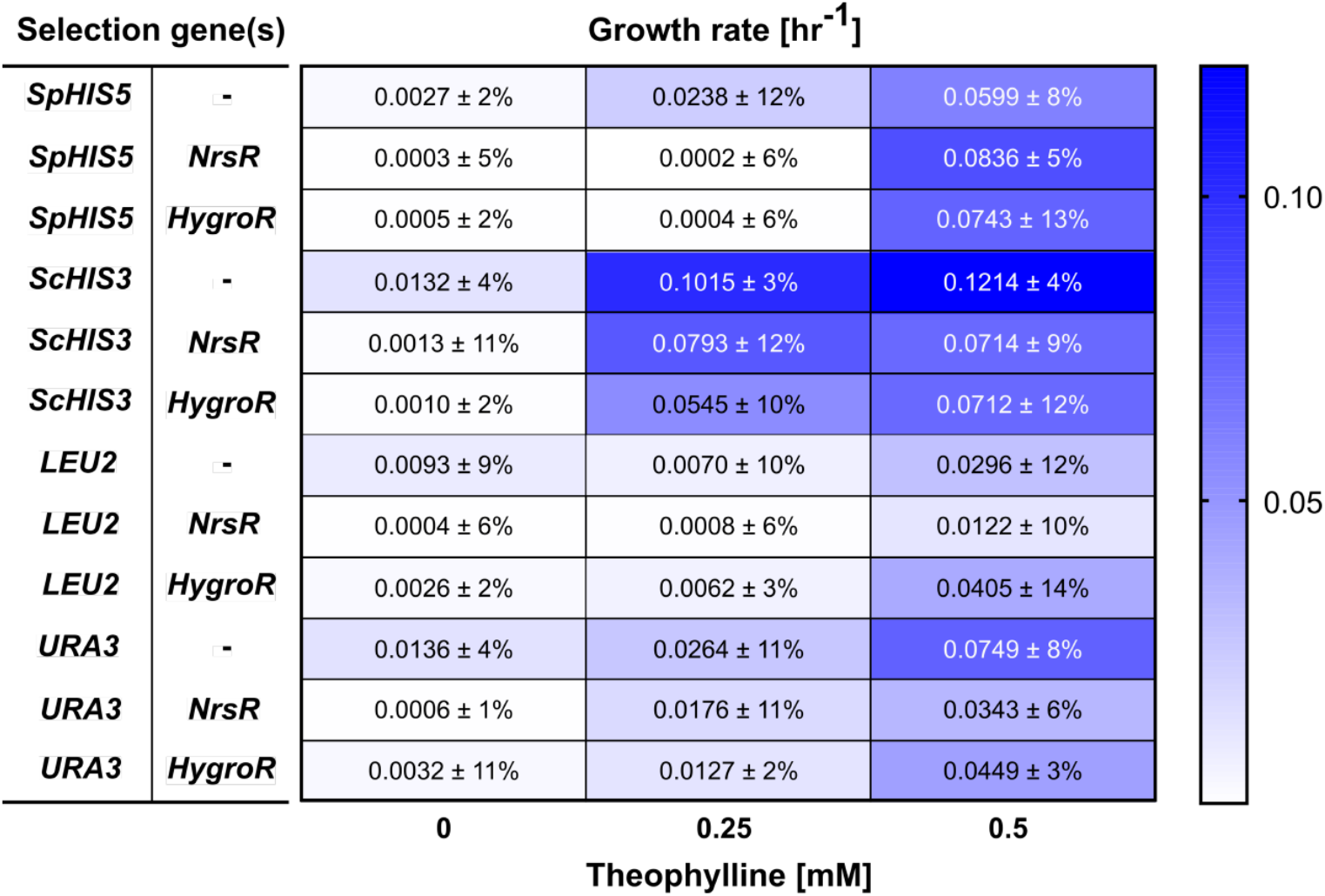
Growth rate measured from selection strains incorporating both an auxotrophic gene and an antibiotic resistance gene, titrated by a theophylline-responsive RNA switch. For each selection strain, an auxotrophic gene (*SpHIS5, ScHIS3, LEU2*, or *URA3*) is paired with an antibiotic resistance gene (*NrsR* or *HygroR*), and both selection genes are under the regulation of tZ_3_EV6p. Yeast growth rate of each selection strain is calculated from OD600 measured over a time course, induced by 0, 0.25, or 0.5 mM theophylline.

## CONCLUSIONS AND DISCUSSION

We designed and tested an RNA switch-based selection system for yeast growth regulation. We first demonstrated the use of a transcription factor-based gene activation system to enhance the dynamic range of an RNA switch. Using a theophylline-responsive RNA switch as an example, we successfully demonstrated its ability to titrate the activity of auxotrophic genes, i.e., *URA3, SpHIS5*, and *LEU2*, for yeast growth regulation. We further validated the use of *Sp*HIS5 for enriching a high-producing variant of CDM among a random mutagenesis library of low-producing variants of CDM. Specifically, we demonstrated enrichment of a high-producing CDM variant by 200-fold from 0.1% initial presence to a final 20% of the population. We next demonstrated the ability to incorporate a second selection pressure, i.e., antibiotics, into the selection system to distinguish growth among low-producing variants. We identified antibiotic resistance genes *NrsR* and *HygroR* as two candidates for cooperation with auxotrophy-based selection. Via fed theophylline assays, we demonstrated that expression of both *NrsR* and *HygroR* can eliminate growth of mid-to-low level variants whereas allowing growth of mid-to-high level variants of CDM.

## METHOD AND MATERIALS

### Plasmid construction

All the primers in this work were synthesized by the Stanford Protein and Nucleic Acid Facility (Stanford, CA). PCR amplifications were performed with Q5 High-Fidelity DNA polymerase (NEB, Ipswich, MA), and PCR products were purified using the DNA Clean and Concentrator Kit (Zymo Research, Irvine, CA). Plasmids generated in this work are listed in Table 1.

**Table 1:**
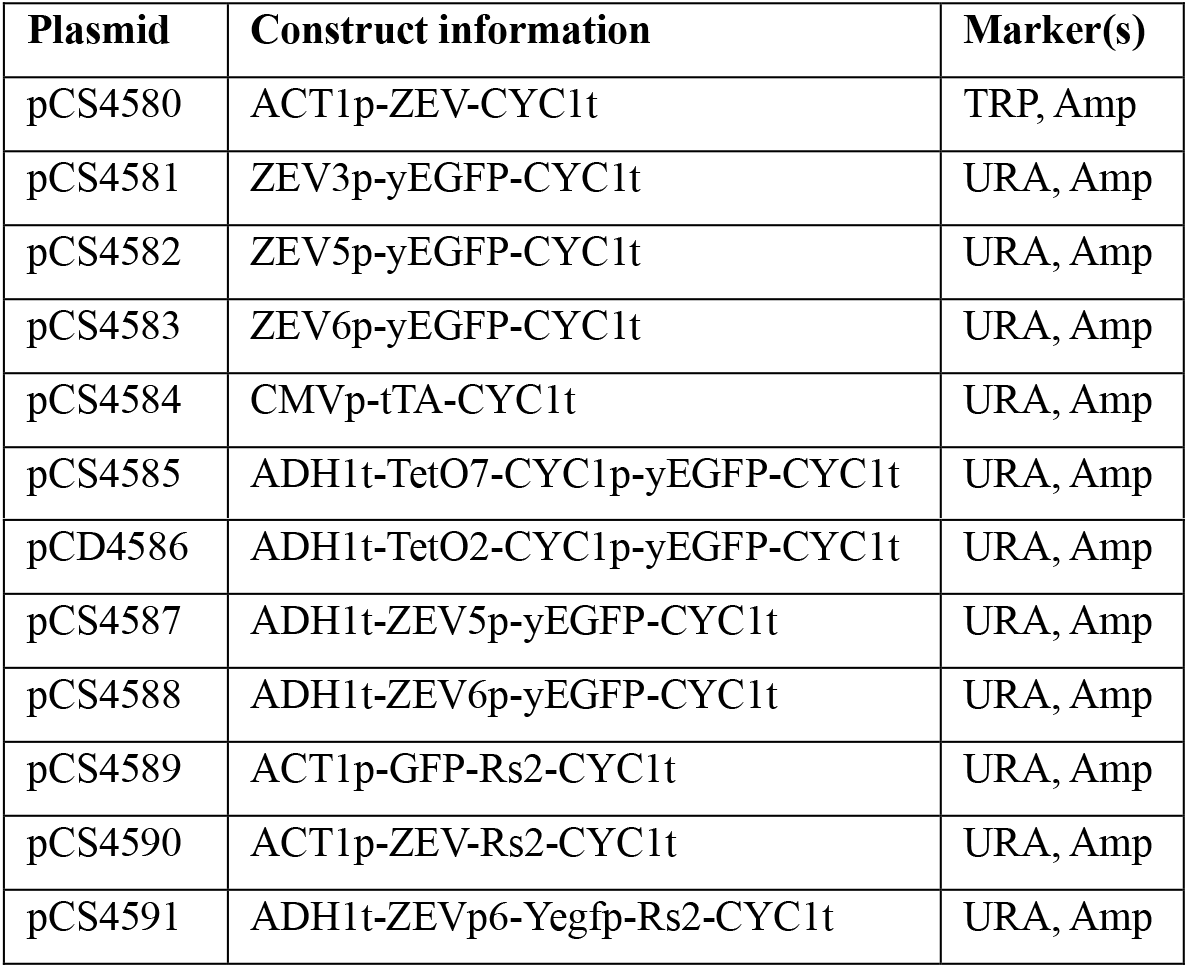
Plasmid constructed for RNA switch-based selection system.

| Plasmid | Construct information | Marker(s) |
| --- | --- | --- |
| pCS4580 | ACT1p-ZEV-CYC1t | TRP, Amp |
| pCS4581 | ZEV3p-yEGFP-CYC1t | URA, Amp |
| pCS4582 | ZEV5p-yEGFP-CYC1t | URA, Amp |
| pCS4583 | ZEV6p-yEGFP-CYC1t | URA, Amp |
| pCS4584 | CMVp-tTA-CYC1t | URA, Amp |
| pCS4585 | ADH1t-TetO7-CYC1p-yEGFP-CYC1t | URA, Amp |
| pCD4586 | ADH1t-TetO2-CYC1p-yEGFP-CYC1t | URA, Amp |
| pCS4587 | ADH1t-ZEV5p-yEGFP-CYC1t | URA, Amp |
| pCS4588 | ADH1t-ZEV6p-yEGFP-CYC1t | URA, Amp |
| pCS4589 | ACT1p-GFP-Rs2-CYC1t | URA, Amp |
| pCS4590 | ACT1p-ZEV-Rs2-CYC1t | URA, Amp |
| pCS4591 | ADH1t-ZEVp6-Yegfp-Rs2-CYC1t | URA, Amp |

### Yeast strain construction and transformation

Yeast strains used in this study are listed in Table 2. All yeast strains are haploid, derived from CEN.PK2-1D (MATα URA3-52, TRP1-289, LEU2-3/112, HIS3D1, MAL2-8C, and SUC2), referred to as CEN.PK2. For yeast transformations, a single colony of the parent strain was inoculated in yeast peptone with 2% dextrose (YPD) media and incubated overnight at 30°C and 220 rpm. The saturated overnight culture was then diluted 50-fold in fresh YPD media and incubated for 4 to 6 hours. A volume of 2.5 ml yeast cells was used per transformation. The cells were then harvested by centrifugation at 3500g for 4 min and prepared for transformation using the Frozen-EZ Yeast Transformation II Kit (Zymo Research, Irvine, CA). For plasmid transformations, 50 ng of DNA was used per transformation. The transformed cells were plated directly onto synthetic dropout agar plates after 45-min incubation with EZ3 solution. For Cas9-based chromosomal integrations, 100 ng of the Cas9 plasmid (encodes G418 resistance) and 500 ng of the linear DNA fragments were used per transformation, and the transformed cells were subject to a 2-hour recovery at 30°C in YPD media after 45-min incubation with EZ3 solution. The cells were plated onto synthetic dropout plates supplemented with G418 (400 mg/liter) to select for colonies with successfully integrated constructs. The plate cultures were incubated 2 to 3 days before colonies were picked for growth assays.

**Table 2:** Strain constructed for RNA switch-based selection system.

| Strain | Genotype |
| --- | --- |
| CEN.PK2-1D | MATα ura3-52, trp1-289; leu2-3/112, his3Δ1, MAL2-8C, SUC2 |
| CSY1362 | CEN.PK2-1D; *LYP1*:: ACT1p-yEGFP-Rs2-CYC1t |
| CSY1363 | CEN.PK2-1D; *URA3*:: ACT1p-Z<sub>3</sub>EV-Rs2-CYC1t |
| CSY1364 | yDKzev2; *LYP1*:: ADH1t-Z<sub>3</sub>EV6p-yEGFP-Rs2-CYC1t |
| CSY1365 | yDKzev2; *LYP1*:: ADH1t-Z<sub>3</sub>EV6p-yEGFP-CYC1t |
| CSY1366 | CEN.PK2-1D; *LYP1*:: tZ<sub>3</sub>EV6p-yEGFP-CYC1t |
| CSY1367 | yDKzev9; *URA3*:: CYC1p-Z<sub>3</sub>EV-Rs2-CYC1t |
| CSY1368 | yDKzev9; *URA3*:: PYKp-Z<sub>3</sub>EV-Rs2-CYC1t |
| CSY1369 | yDKzev9; *URA3*:: TEFp-Z<sub>3</sub>EV-Rs2-CYC1t |
| CSY1370 | yDKzev9; *URA3*:: GPDp-Z<sub>3</sub>EV-Rs2-CYC1t |
| CSY1371 | CEN.PK2-1D; *URA3*:: TEFp-Z<sub>3</sub>EV-Rs2-CYC1t |
| CSY1372 | yDKzev16; *LYP1*:: ADH1t-Z<sub>3</sub>EV6p-URA3-CYC1t |
| CSY1373 | yDKzev16; *LYP1*:: ADH1t-Z<sub>3</sub>EV6p-*Sp*HIS5-CYC1t |
| CSY1374 | yDKzev16; *LYP1*:: ADH1t-Z<sub>3</sub>EV6p-TRP1-CYC1t |
| CSY1375 | yDKzev16; *LYP1*:: ADH1t-Z<sub>3</sub>EV6p-LEU2-CYC1t |
| CSY1376 | yDKzev16; *LYP1*:: ADH1t-Z<sub>3</sub>EV6p-KanR-CYC1t |
| CSY1377 | yDKzev16; *LYP1*:: ADH1t-Z<sub>3</sub>EV6p-HygroR-CYC1t |
| CSY1378 | yDKzev16; *LYP1*:: ADH1t-Z<sub>3</sub>EV6p-NrsR-CYC1t |
| CSY1379 | yDKzev16; *LYP1*:: ADH1t-Z<sub>3</sub>EV6p-CyhR-CYC1t |
| CSY1380 | yDKsele11; *HIS2*:: ADH1t-Z<sub>3</sub>EV6p-NrsR-CYC1t |
| CSY1381 | yDKzev16; *LYP1*:: ADH1t-Z<sub>3</sub>EV6p-HygroR-CYC1t |
| CSY1382 | yDKsele33; *HIS2*:: ADH1t-Z<sub>3</sub>EV6p-*Sc*HIS3-CYC1t |
| CSY1383 | yDKsele32; *HIS2*:: ADH1t-Z<sub>3</sub>EV6p-HygroR-CYC1t |
| CSY1384 | yDKsele15; *HIS2*:: ADH1t-Z<sub>3</sub>EV6p-*Sc*HIS3-CYC1t |
| CSY1385 | yDKsele33; *HIS2*:: ADH1t-Z<sub>3</sub>EV6p-LEU2-CYC1t |
| CSY1386 | yDKsele13; *HIS2*:: ADH1t-Z<sub>3</sub>EV6p-NrsR-CYC1t |
| CSY1387 | yDKsele15; *HIS2*:: ADH1t-Z<sub>3</sub>EV6p-LEU2-CYC1t |
| CSY1388 | yDKsele33; *HIS2*:: ADH1t-Z<sub>3</sub>EV6p-URA3-CYC1t |
| CSY1389 | yDKsele15; *HIS2*:: ADH1t-Z<sub>3</sub>EV6p-URA3-CYC1t |
| CSY1390 | yDKzev16; *LYP1*:: ADH1t-Z<sub>3</sub>EV6p-*Sc*HIS3-CYC1t |

### Yeast growth measurements

Yeast strains were inoculated in triplicates into a 400 μl culture of synthetic complete medium or synthetic dropout medium in 96-well plates and grown overnight to saturation prior to 100-fold back dilution into a 400 μl culture of fresh medium supplemented with 500 nM β-estradiol. OD600 of the culture was then measured at 0, 6, 12, 24, and / or 56 hours of growth using Tecan Safire2 plate reader (10s orbital shaking prior to measurement, final absorbance collected from an average of 15 laser readings). For each time point, 100 ul of culture was sampled and transferred back to the culture plate for further growth after OD600 measurement. The mean and standard deviation of OD600 was then calculated from the triplicates of each strain at each time point. For calculating growth rates from OD600 measurements, the following equation was used: yeast growth rate = ln(OD600_(t=tp2)_ / OD600_(t=tp1)_)/△tp. Time points (tps) were chosen such that the growth curve data between the time points represented exponential growth phase of yeast cells.

### Sample preparation and data analysis for next generation sequencing

Yeast cells transformed with plasmids encoding the CDM library or control variant are harvested and the plasmid DNA extracted using the Zymoprep Yeast Plasmid Miniprep I Kit (Zymo Research, Irvine, CA). DNA sequences of the variant library are then PCR-amplified using primers with NGS adapters and gel-extracted from the reaction mixture by length of the PCR product. The samples were sequenced on MiSeq machines at 2 × 150 bp. NGS data were cleaned up by adapter trimming and read quality was controlled such that any reads with a Q-score < 30 and a length < 250 bp will be removed. The reads were then aligned to CDM sequence template using Bowtie2 end2end alignment and the alignment data visualized using R.

## Notes

### Competing Interest Statement

The authors have declared no competing interest.

