## Supplementary Information for "Building an RNA switch-based selection system for enzyme evolution in yeast"

**SUPPLEMENTARY DOCUMENTS**

**Mathematical modelling of RNA switch-based regulation for gene activation.** An ordinary differential equation (ODE)-based model was built to capture system behavior, i.e., EGFP expression, under RNA switch-based regulation for each of the constructs tested in yeast, as shown in Table 3. The modelling of transcription and translation processes, transcription factor-based gene activation, and RNA switch-based regulation is shown in Table 4.

| System construct | System ODEs |
|---|---|
| i. 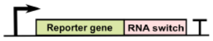  | $\frac{d[m_{GFP}]}{dt} = k_{trsTEF} - (fxn_{RNA1}([L])k_{mloss-clv} + fxn_{RNA2}([L])k_{mloss})[m_{GFP}]$<br>$\frac{d[P_{GFP}]}{dt} = k_{trl}[m_{GFP}] - k_{ploss}[P_{GFP}]$ |
| ii. 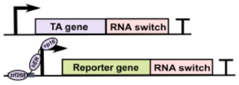  | $\frac{d[m_{TA}]}{dt} = k_{trsTEF} - (fxn_{RNA1}([L])k_{mloss-clv} + fxn_{RNA2}([L])k_{mloss})[m_{TA}]$<br>$\frac{d[P_{TA}]}{dt} = k_{trl}[m_{TA}] - k_{ploss}[P_{TA}]$<br>$\frac{d[m_{GFP}]}{dt} = fxn_{IP}([P_{TA}][I])\ k_{trsZEV} - (fxn_{RNA1}([L])k_{mloss-clv} + fxn_{RNA2}([L])k_{mloss})[m_{GFP}]$<br>$\frac{d[P_{GFP}]}{dt} = k_{trl}[m_{GFP}] - k_{ploss}[P_{GFP}]$ |
| iii. 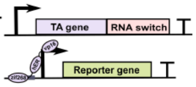  | $\frac{d[m_{TA}]}{dt} = k_{trsTEF} - (fxn_{RNA1}([L])k_{mloss-clv} + fxn_{RNA2}([L])k_{mloss})[m_{TA}]$<br>$\frac{d[P_{TA}]}{dt} = k_{trl}[m_{TA}] - k_{ploss}[P_{TA}]$<br>$\frac{d[m_{GFP}]}{dt} = fxn_{IP}([P_{TA}][I])\ k_{trsZEV} - k_{mloss}[m_{GFP}]$<br>$\frac{d[P_{GFP}]}{dt} = k_{trl}[m_{GFP}] - k_{ploss}[P_{GFP}]$ |

**Supplementary table 1:** ODE-based modelling of gene expression regulated by RNA switch-based constructs tested in yeast.

Basic transcription/translation model:

$$\frac{d[m]}{dt} = k_{trs} - k_{mloss}[m]$$

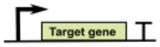


$$\frac{d[P]}{dt} = k_{trl}[m] - k_{ploss}[P]$$

Inducible promoter-regulated transcription model:

$$\frac{d[m]}{dt} = \mathbf{fxn_{IP}([P_{TA}][I])}k_{trs} - k_{mloss}[m]$$

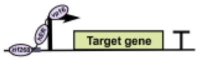


RNA switch-regulated transcription model:

$$\frac{d[m]}{dt} = k_{trs} - \mathbf{fxn_{RNA1}}([L])k_{mloss-clv}[m] - \mathbf{fxn_{RNA2}}([L])k_{mloss}[m]$$

$Z_3EVp$ induction function:

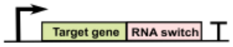


$$P_{TA} + I \xrightarrow{K_I} PI$$

$$PI + DNA \xrightarrow{K_{PI}} PI - DNA$$

$$\mathbf{fxn_{IP}([P_{TA}][I])} = \frac{K_{PI}K_I[I][P_{TA}]^n}{1 + K_{PI}K_I[I][P_{TA}]^n}$$

RNA switch regulation function:

$$\mathbf{fxn_{RNA1}}([L]) = \frac{p}{1 + K_L[L]}$$

$$\mathbf{fxn_{RNA2}}([L]) = \frac{1 + K_L[L] - p}{1 + K_L[L]}$$

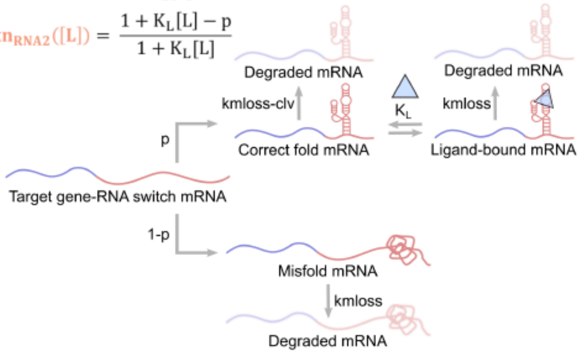


Species definitions:

m: mRNA

P: protein

$P_{TA}$: TA protein

I: Inducer (β-estradiol)

PI: Induced TA protein

PI-DNA: TA protein-bound $Z_3EVp$

L: ligand

Parameters:

$k_{trs}$: transcription rate

$k_{mloss}$: mRNA degradation rate

$k_{trl}$: translation rate

$k_{ploss}$: protein degradation rate

$k_{mloss\text{-}clv}$: cleaved RNA switch degradation rate

$K_I$: association constant of $P_{TA}$ and I

$K_{PI}$: association constant of $P_I$ and DNA

$K_L$: association constant of correct fold mRNA and the ligand

n: Hill coefficient for modelling cooperativity in $Z_3EV$ protein binding

p: proportion of correct fold mRNA

**Supplementary table 2:** Mathematical modelling of transcriptional and post-transcriptional regulation by genetic components in the ODE system.
